# Epstein-Barr Virus Encoded lncRNAs Control the Viral Lytic Switch

**DOI:** 10.1101/2025.10.29.685446

**Authors:** Zhixuan Li, Yifei Liao, Weiyue Ding, Davide Maestri, Scott L Grote, Gabriel A Romero Agosto, Laura A Murray-Nerger, Rui Guo, Bo Zhao, Silvi Rouskin, Mingxiang Teng, Benjamin E Gewurz

## Abstract

Epstein-Barr virus (EBV) uses a biphasic lifecycle, switching between latent and lytic phases to persistently infect most adults. Latency is observed in most tumor cells of the 200,000 EBV-associated cancers/year. EBV reactivation is increasingly implicated in autoimmune diseases, including multiple sclerosis. However, mechanisms that regulate EBV reactivation have remained incompletely understood. Here, we leveraged multi-omic approaches to reveal the existence of pro-latency and a pro-lytic viral long noncoding RNAs (lncRNAs) that counter-regulate the lytic switch. Known reactivation triggers rapidly induced expression of the pro-lytic lncRNA, which encodes an RNA G-quadruplex that mediated its interaction with CTCF. The pro-lytic lncRNA occupies viral origin of lytic replication enhancers and promotes their looping to the immediate early lytic promoter to trigger reactivation. The pro-latency lncRNA duplexes with the pro-lytic RNA to impede its interactions with CTCF. These studies lay a foundation for therapeutic approaches to manipulate the EBV lytic switch.

## Introduction

Epstein-Barr virus (EBV) persistently infects ∼95% of adults worldwide. It is associated with >200,000 cancer cases per year, including Burkitt, Hodgkin and diffuse large B-cell lymphomas, nasopharyngeal and gastric carcinomas^1-4^. EBV also causes post-transplant lymphoproliferative diseases, T and natural killer cell lymphomas^5,6^. Together, these comprise nearly 1% of all human cancers. EBV causes infectious mononucleosis and is increasingly linked to a range of autoimmune diseases, including multiple sclerosis, systemic lupus erythematosus, rheumatoid arthritis and primary biliary cholangitis, where lytic reactivation may be a major trigger^1,7,8^.

Upon B-cell infection, the double stranded DNA EBV genome is rapidly chromatinized and circularized. Epigenetic mechanisms silence expression of most of the ∼80 EBV-encoded genes, the majority of which remain repressed until EBV undergoes lytic reactivation. EBV uses a series of latency programs, in which combinations of 1-8 viral latency oncogenes and non-coding RNA are expressed, to drive infected B cell proliferation and differentiation into memory cells, the reservoir of lifelong EBV infection^9-12^. Within memory cells, EBV uses the latency I program, in which EBNA1 is the only viral protein expressed together with EBV miRNAs and EBER non-coding RNAs, and even EBNA1 can be silenced in resting memory cells^13^. Burkitt lymphoma typically use the latency I program to evade antiviral cell mediated immune responses. These observations suggest that host factors and perhaps also viral non-coding RNAs mediate EBV B-cell latency.

To define host factors critical for EBV latency in B-cells, we performed human genome-wide CRISPR-Cas9 screens. These identified that the proto-oncogene c-MYC, which is highly expressed in Burkitt lymphoma tumors, as well as LSD1, CoREST and ZNF217^14,15^, are each necessary for maintenance of EBV latency. Depletion of any of these, of histone chaperones or histone H1 or factors critical for maintenance of DNA methylation^16-18^ triggered EBV reactivation by incompletely understood mechanisms.

Reactivation stimuli such as by plasma membrane immunoglobulin cross-linking induce two EBV immediate early genes that encode the BZLF1 and BRLF1 transcription factors^12,19^. Each are critical for completion of the B cell EBV lytic cycle^20^. However, the ability of BZLF1 versus BRLF1 to initiate reactivation varies by cell type^21^, with BZLF1 playing a more important role in cells with CpG hypermethylation, including Burkitt lymphoma cells^22-24^. EBV immediate early proteins induce expression of ∼35 viral early genes, whose gene-products mediate replication of viral DNA and include a kinase that activates the cytotoxic activity of the antiviral ganciclovir^25^. Unchromatinized, linear EBV genomes produced by the lytic cycle are the template for the expression of ∼35 viral late genes, which encode structural proteins and factors necessary for virion assembly and release^26^. The lytic cycle amplifies EBV genomes by 10-1000-fold^27^.

While *oriLyt* enhancers are critical drivers of late lytic gene expression^28,29^ and of viral lytic DNA synthesis, we recently suggested that they are also critical drivers of EBV lytic reactivation^14^. Interestingly, *oriLyt* encode multiple transcripts produced even in viral latency^30^, the function of which remain to be fully elucidated. Depletion of MYC, the histone lysine demethylase LSD1 or immunoglobulin cross-linking rapidly induce long-range DNA interactions between *oriLyt* and the immediate early *BZLF1* promoter that drive the B cell lytic cycle. Disruption of MYC-occupied *oriLyt* E-box sites also drives reactivation^14,15^. Together, these suggest that a complex containing MYC and LSD1 may serve to promote viral latency by controlling EBV genomic higher order architecture by incompletely understood mechanisms.

Most available therapies do not harness the presence of EBV genomes within tumor cells, particularly in the highly restricted latency I program. Consequently, there is growing interest in strategies that harness EBV lytic reactivation together with ganciclovir, whose cytotoxicity is licensed by the EBV lytic kinase BGLF4^25,31^. There is also increasing evidence that the viral lytic cycle contributes to lymphomagenesis and to the tumor microenvironment of epithelial tumors^32-36^.

Here, we integrated complimentary approaches to investigate key mechanisms underlying EBV genomic higher order architecture and its relationship with viral reactivation. HiChIP^37^ further highlighted *oriLyt:BZLF1* long-range interactions upon EBV reactivation. Degron-based^38^ depletion of RAD21 cohesin, which is a key driver of DNA loop formation, strongly impaired EBV reactivation. Mechanistically, we identified pro-latency and pro-lytic lncRNAs encoded by EBV *oriLyt*, the expression of latter of which was repressed by MYC. oriLyt pro-lytic lncRNA expression was rapidly induced upon treatment with lytic induction stimuli, and its knockdown impaired reactivation. *oriLyt* lncRNA bound to the DNA loop anchor CTCF in a manner dependent on its RNA G4 quadruplex.

## Results

### Immunoglobulin crosslinking drives looping between both origins of lytic replication and the lytic cycle inducing BZLF1 promoter

To define how EBV genomic architecture remodels upon lytic reactivation, EBV+ Akata Burkitt B cells were mock induced, or induced for lytic reactivation by αIgG crosslinking for 24 hours. We then utilized histone 3 lysine 27 acetyl (H3K27Ac) HiChIP^37^ to quantitate long-range interactions between EBV genomic sites marked by H3K27Ac, deposited at active enhancers and promoters (**Figure 1A**). This analysis highlighted that new DNA contacts formed between each *oriLyt* enhancer and the *BZLF1* immediate early gene region (**Figure 1B-C, S1A**). αIgG cross-linking also induced contacts between the terminal repeat region and the *BZLF1* promoter, whereas a DNA loop between the *BZLF1* promoter and *BILF2* region was lost (**Figure 1B-C, S1A**). Taken together with chromatin conformation capture (3C) analyses^14,15^, these results indicate that EBV higher order architecture remodels with reactivation to juxtapose *oriLyt* enhancers with the immediate early *BZLF1* promoter.

**Figure 1.**
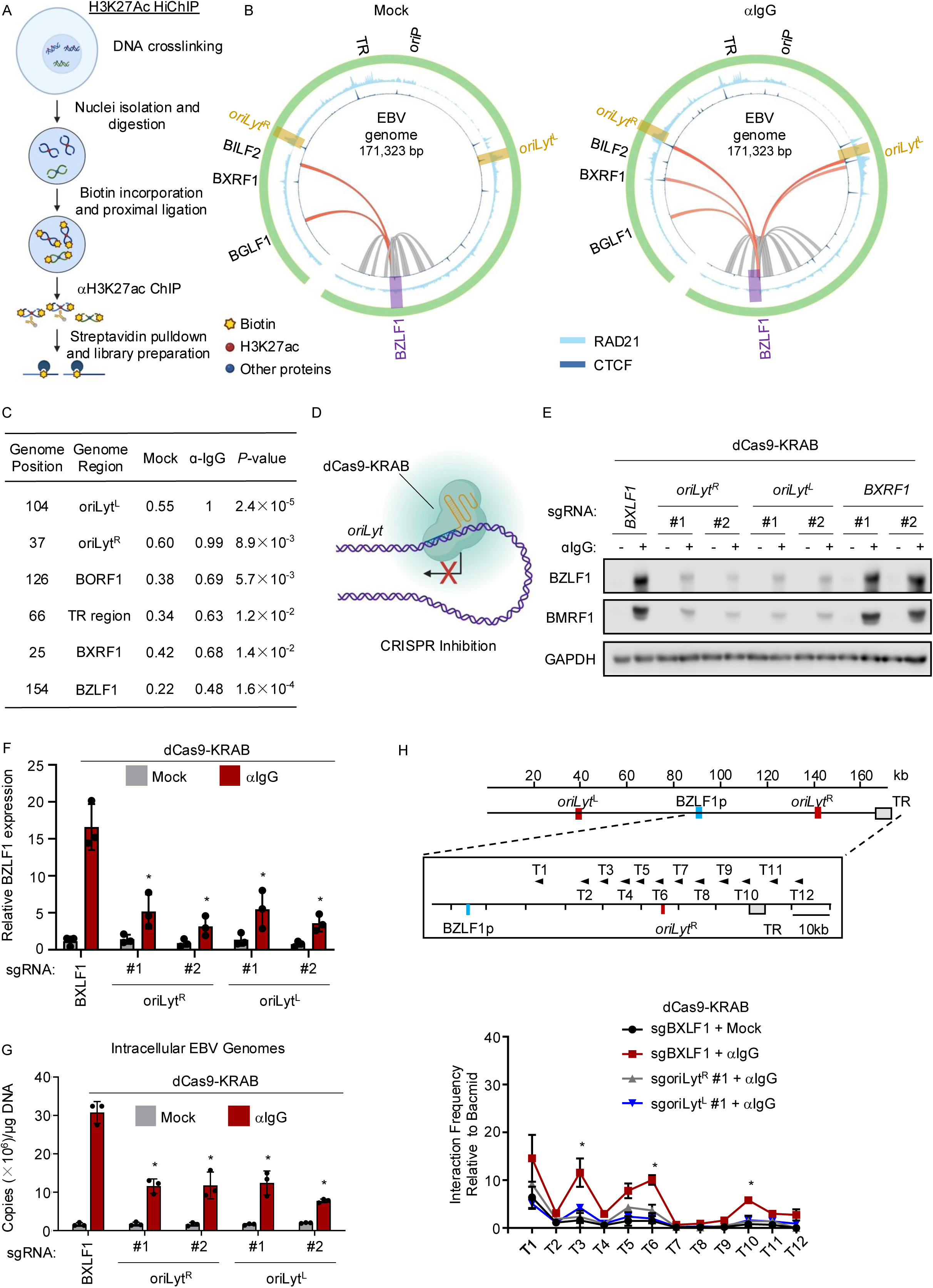
Ig crosslinking triggered *oriLyt and BZLF1* promoter association, which is important for EBV lytic reactivation. (A) Schematic of H3K27ac HiChIP assays. Following Akata Burkitt cell formaldehyde crosslinking, nuclei were isolated and digested by MboI. Sheared ends were biotinylated and ligated. Histone3 lysine 27 acetyl (H3K27ac) activated chromatin immunoprecipitation was followed by biotinylated DNA streptavidin pulldown. (B) Circos plots of EBV genomic H3K27Ac HiChIP from Akata cells mock induced (left) or induced for EBV reactivation by αIgG crosslinking for 24 hours (10μg/ml) (right). Gray and orange arcs indicate short range interactions (<20kb). Long range interactions (>20kb) are depicted by red and orange arcs. Data are from n=2 independent replicates. Genomic positions of selected EBV genes, the origins of lytic replication (*oriLyt^R^*and *oriLyt^L^*), the origin of plasmid replication (*oriP*) and the terminal repeats (TR) are indicated. Also shown are Akata RAD21 (orange) and CTCF (blue) ChIP-seq tracks from n=2 replicates. (C) Regions with the most enriched EBV-genomic long-range DNA interactions following αIgG crosslinking. Shown are EBV genome positions and annotation at regions with increased interaction frequency, interaction scores in αIgG versus mock-stimulated Akata cells. (D) Schematic diagram of CRISPR interference at *oriLyt* enhancer regions. (E) Immunoblot of whole cell lysates (WCL) from Akata cells stably expressing dCas9-KRAB and sgRNAs targeting the indicated EBV genomic regions, unstimulated or 24 hours post-αIgG crosslinking. Blots are representative of n=3 replicates. (F) Mean ± S.D. of BZLF1 mRNA expression levels from n=3 replicates of dCas9-KRAB Akata cells expressing the indicated sgRNAs and mock-induced or induced for EBV lytic reactivation by αIgG crosslinking for 24 hours. sgRNA targeting BXLF1 was used as control. (G) Mean ± S.D. of EBV intracellular genome copy number from n=3 replicates of dCas9-KRAB+ Akata cells expressing the indicated sgRNAs and mock induced or induced for EBV reactivation by αIgG crosslinking for 24 hours. P-values compare the indicated to αIgG BXLF1 control values. (H) Chromatin conformation capture (3C) analysis of BZLF1 promoter interaction frequency with the indicated EBV genomic regions. 3C was performed with BZLF1 promoter anchor primer (blue rectangle) and the indicated test (T) primers in dCas9-KRAB+ Akata cells expressing the indicated sgRNAs and mock induced or induced by αIgG crosslinking. sgRNA targeting BXLF1 was used as control. Shown are mean ± S.D. 3C assay interaction frequencies from 3 replicates, normalized by EBV bacmid input values. P-values compare the indicated to αIgG BXLF1 control values. *p < 0.05

We then utilized CRISPR interference (CRISPRi), in which a single guide RNA (sgRNA) targets a catalytically dead *Streptococcus pyogenes* Cas9/KRAB transcription repressor fusion^39^ to a specified genomic site (**Figure 1D**), to test *oriLyt* enhancer roles in EBV reactivation. We expressed sgRNAs targeting *oriLyt R* or *L* or control EBV genomic *BXLF1* or *BXRF1* regions. CRISPR interference of either *oriLyt* enhancer using published sgRNAs^40^ strongly impaired EBV reactivation in response to αIg crosslinking, as judged by BZLF1 and BMRF1 early protein induction, lytic EBV genome amplification or secretion of encapsidated EBV DNA (**Figure 1E-G** and **S1B-C**). By contrast, CRISPRi targeting of the BILF2 region that loops to BZLF1 in latency did not appreciably alter EBV lytic protein expression in Mutu I triggered for reactivation by αIgM cross-linking (**Figure S1D**).

CRISPRi against either *oriLyt*, but not against BXLF1, similarly impaired BZLF1 and BMRF1 induction by protein kinase C agonist 12-O-Tetradecanoylphorbol-13-acetate (TPA, also called phorbol myristate acetate) or histone deacetylase inhibitor sodium butyrate (NaB) (**Fig. S1E**). CRISPRi against either *oriLyt* also impaired Ig cross-linking or TPA/NaB induced long-range *oriLyt*:*BZLF1* promoter region interactions, as well as to a somewhat lesser extent interactions between EBV terminal repeats (TR) and the BZLF1 promoter (**Figure 1H and S1F**). Taken together, these results indicate that immunoglobulin crosslinking results in long-range interactions between *oriLyt* and BZLF1 promoter activated chromatin, and that this is critical for EBV B-cell reactivation.

### DNA looping factor cohesin and CTCF roles in EBV reactivation

Long-range DNA loops are formed by the activity of cohesins, which form DNA-entrapping rings comprised of structural maintenance complex (SMC) proteins, RAD21 and stromal antigen (STAG) 1 or 2^41,42^. Cohesins use ATPase activity to extrude DNA at rates that can approach 0.5-2 kilobase of DNA per second^43,44^. Cohesins extrude DNA until they encounter CTCF bound to convergent CTCF binding sites (CBS)^45-47^. Notably, the *BZLF1* promoter and *oriLyt* regions contain convergent CTCF binding sites. We performed ChIP-seq to map EBV genomic RAD21 cohesin and CTCF occupancy in latency and upon Akata B cell EBV reactivation. αIgG crosslinking rapidly increased RAD21 and CTCF occupancy at both *oriLyt* and to a lesser extent at the *BZLF1* promoter (**Figure 2A**). EBV genomic RAD21 and CTCF ChIP-seq signals were highest at *oriLyt* in reactivated cells, further suggesting *oriLyt* DNA looping roles. Increased *oriLyt* and *BZLF1* promoter CTCF and RAD21 occupancy were also observed in TPA and NaB treated Akata cells by ChIP-qPCR analysis (**Figure S2A**).

**Figure 2.**
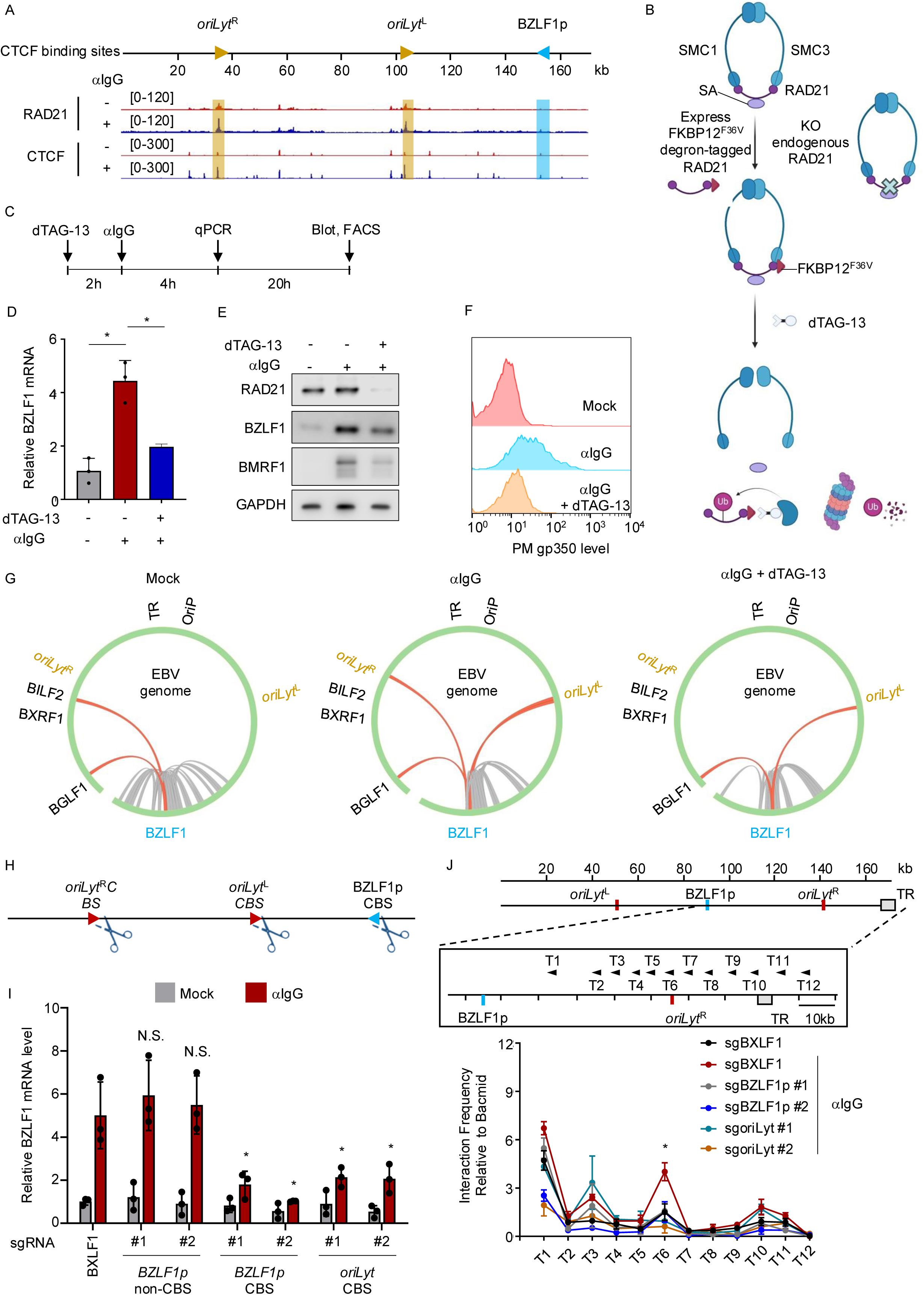
RAD21 cohesin is critical for *oriLyt/BZLF1* promoter looping and EBV reactivation. (A) ChIP-seq analysis of EBV genomic CTCF and RAD21 occupancy. ChIP-seq RAD21 (top) and CTCF (bottom) tracks from Akata cells mock stimulated or stimulated by αIgG for 4 hours. Average values from 2 independent replicates. *OriLyt* and *BZLF1* promoter (BZLF1p) CBS orientations are shown. (B) Construction of Akata cells with degron-tagged RAD21. FKBP12^F36V^-degron tagged RAD21 was overexpressed, and endogenous RAD21 was knocked out by CRISPR. Degron-tagged RAD21 can be rapidly depleted by addition of dTAG-13. (C) Experimental plan schematic diagram. (D) Mean ± S.D. BZLF1 mRNA levels from 3 qPCR replicates of Akata cells mock treated or treated with dTAG13 (100nM) for two hours prior to αIgG for 4 hours. (E) Immunoblot of WCL from Akata cells treated with dTAG-13 for 2 hours to deplete degron-tagged RAD21 and then αIgG for 24 hours as indicated. Blots are representative of 3 replicates. (F) Flow cytometry (FACS) analysis of plasma membrane (PM) gp350 expression in Akata cells mock treated or treated with dTAG-13 for 2 hours and then by αIgG for 24 hours, as indicated. Representative of 3 replicates. (G) EBV genomic H3K27Ac HiChIP circos plots from Akata cells, mock induced (left) or induced by αIgG for 24 hours in the absence (middle) or 2 hours post-dTAG (100nM, right). Red arcs, >20kb DNA interactions between H3K27ac marked regions from 2 replicates. (H) CRISPR editing of *oriLyt* vs *BZLF1* promoter CBS. (I) qPCR mean ± S.D. BZLF1 mRNA levels from 3 replicates of Cas9+ Akata cells expressing the indicated sgRNAs against EBV genomic controls (*BXLF1* or *BZLF1p* sites not containing a CBS) vs sgRNAs targeting *BZLF1p* or *oriLyt* CBS sites and mock treated or treated with αIgG for 24 hours. P-values cross-compare the indicated with control BXLF1 αIgG values. (J) 3C analysis of *BZLF1* promoter interaction frequency with the indicated EBV genomic regions. 3C was performed with *BZLF1p* anchor primer (blue rectangle) and the indicated test (T) primers in Cas9+ Akata cells expressing the indicated sgRNAs and mock induced or induced by αIgG. Shown are mean ± S.D. 3C assay interaction frequencies from 3 replicates, normalized by EBV bacmid input values. *p < 0.05

We next tested whether DNA looping was necessary for Burkitt EBV reactivation. To more rapidly and completely deplete RAD21 cohesin levels than is achievable by CRISPR editing, where CRISPR knockout does not deplete pre-loaded DNA-bound cohesin complexes, we engineered a system for rapid protein-level RAD21 depletion. First, we stably expressed a C-terminally FKBP12^F36V^-tagged RAD21 allele at near physiological levels in Akata cells. Addition of the heterobifunctional degrader dTAG13^38^ rapidly targets FKBP12^F36V^-tagged proteins for proteasomal degradation (**Figure 2B**). Next, we knocked out endogenous RAD21, such that only the FKBP12^F36V^-tagged RAD21 allele was expressed (**Figure S2B**). dTAG-13 treatment rapidly degraded FKBP12^F36V^-tagged RAD21 to nearly undetectable levels within 2 hours. Although RAD21 depletion also mildly reduced MYC abundance, likely because DNA looping between immunoglobulin enhancers and the *MYC* promoter contributes to constitutive MYC expression in many Burkitt tumors^48^, BZLF1 or BMRF1 were not de-repressed by RAD21 depletion for up to 24 hours (**Figure S2C**).

We leveraged this system to ask whether RAD21 and therefore also DNA looping were necessary for EBV reactivation. Following mock treatment or RAD21 depletion by two hours of dTAG-13 treatment, we tested effects of αIgG treatment on EBV reactivation (**Figure 2C**). At 4 hours post-IgG crosslinking, BZLF1 mRNA levels were significantly decreased by RAD21 depletion (**Figures 2D**). Further suggestive of key cohesin roles in EBV lytic reactivation, BZLF1, BMRF1 protein and late lytic gp350 expression were each impaired by RAD21 depletion just prior to αIgG crosslinking (**Figures 2E-F**). Importantly, doxycycline-induced BZLF1 cDNA expression bypassed RAD21 depletion effects on EBV BMRF1 early gene expression (**Figure S2D**), suggesting that RAD21 withdrawal specifically impaired EBV immediate early protein induction.

We next used H3K27Ac HiChIP to define conditional RAD21 depletion effects on EBV genomic looping following B-cell receptor stimulation. dTAG13 addition from two hours prior to B-cell receptor cross-linking reduced *oriLyt^R^:BZLF1* looping and decreased the frequency of the *oriLyt^L^:BZLF1* loop (**Figure 2G**). We suspect that a low level of residual DNA-loaded cohesin accounted for residual loop formation in dTAG13 treated cells, though it remains possible that loops form independently of cohesin.

To further test the model that DNA loop extrusion drives B-cell EBV reactivation, we used CRISPR-Cas9 to edit *oriLyt* or *BZLF1* promoter CBS (**Figure 2H**). However, since both *oriLyt* share a CBS sequence, it was not possible to target them individually. As negative controls, we targeted an EBV genomic CBS near the *BXLF1* gene or a non-CBS site within the *BZLF1* promoter. Indel sequencing demonstrated >80% editing efficiency at each of these sites (**Figure S2E**). Importantly, *BZLF1* CBS editing, but not editing of the control *BZLF1* promoter site, significantly reduced *BZLF1* promoter CTCF occupancy (**Figure S2F**). Similarly, *oriLyt* CBS editing decreased CTCF occupancy at *oriLyt*, but not at a control EBV genomic *LMP2A* site (**Figure S2F**).

We then tested EBV genomic CBS editing effects on viral reactivation. CBS editing did not appreciably de-repress lytic gene expression in mock-induced Akata Burkitt cells. However, editing of either the *BZLF1* or *oriLyt* CBS, but not the *BZLF1* promoter control non-CBS site, significantly impaired BZLF1 mRNA upregulation by IgG crosslinking (**Figure 2I**). *BZLF1* promoter or *oriLyt* CBS editing also significantly impaired *BZLF1:oriLyt* looping following αIgG-crosslinking (**Figure 2J**). To address the possibility that *BZLF1* promoter editing might have disrupted its activity, we performed dual luciferase assays on Akata cells that expressed a *BZLF1* promoter driven firefly luciferase reporter. BZLF1 promoter activity was not impaired by CRISPR editing of the *BZLF1* CBS (**Figure S2G**), suggesting that effects on CTCF binding, rather than on the *BZLF1* promoter activity itself, impaired EBV reactivation. Together, these results support a model in which EBV lytic reactivation stimuli trigger juxtaposition of the *oriLyt* enhancers with the *BZLF1* immediate early promoter by CTCF-gated cohesin DNA loop extrusion.

To build on the observation that a RAD21-dependent DNA loop between *BILF2* and *BZLF1* is present in latency, but lost upon lytic reactivation (**Figure 2G**), we next tested the effects of CRISPR editing of the *BILF2* locus CBS. Interestingly, *BILF2* CBS editing by any of three independent sgRNAs de-repressed *BZLF1* mRNA and BZLF1 and BMRF1 protein expression (**Figure S2H-I**). Thus, the BILF2:BZLF1 promoter loop may serve as an insulator to maintain latency in the absence of lytic induction stimuli.

### *oriLyt* encode pro-latency and pro-lytic long non-coding RNAs (lncRNA)

Enhancer activity and transcription often correlate^49-51^. We therefore hypothesized that lytic induction stimuli, such as αIgG crosslinking, rapidly alter *oriLyt* transcription. Either αIgG crosslinking or TPA/NaB treatment rapidly increased *oriLyt* H3K27ac levels, suggestive of chromatin activation (**Figure S3A**). Moreover, αIgG crosslinking increased RNA polymerase II (RNA Pol II) initiation and elongation at *oriLyt*, as shown by ChIP-qPCR analysis of Pol II serine 2 and serine 5 occupancy, respectively (**Figure S3B**).

Enhancer transcription can generate long non-coding RNAs (lncRNA) that modulate transcription factor occupancy, chromatin modification and enhancer-promoter interactions^52-55^. We therefore surveyed for *oriLyt* transcripts produced upon lytic reactivation. First, we re-analyzed strand-specific RNA-seq datasets over the first 24 hours of Akata EBV reactivation by αIgG crosslinking^56^, which identified an *oriLyt* transcript, produced from the EBV genomic positive strand in latency and expressed at higher levels upon reactivation (**Figure 3A**). αIgG treatment also induced a new *oriLyt* negative DNA strand encoded transcript (**Figure 3A**). These transcripts spanned the MYC-bound E-Box site that we previously identified as a major determinant of EBV latency^40^. Consistent with this observation, we also observed transcription from similar EBV genomic regions in published Precision nuclear Run-On sequencing (PRO-seq) analysis of TPA/NaB induced Mutu I Burkitt cells^57^ by 1 hour of treatment (**Figure S3C**). However, PRO-seq analysis failed to unambiguously map *oriLyt* GC-rich repetitive regions.

**Figure 3.**
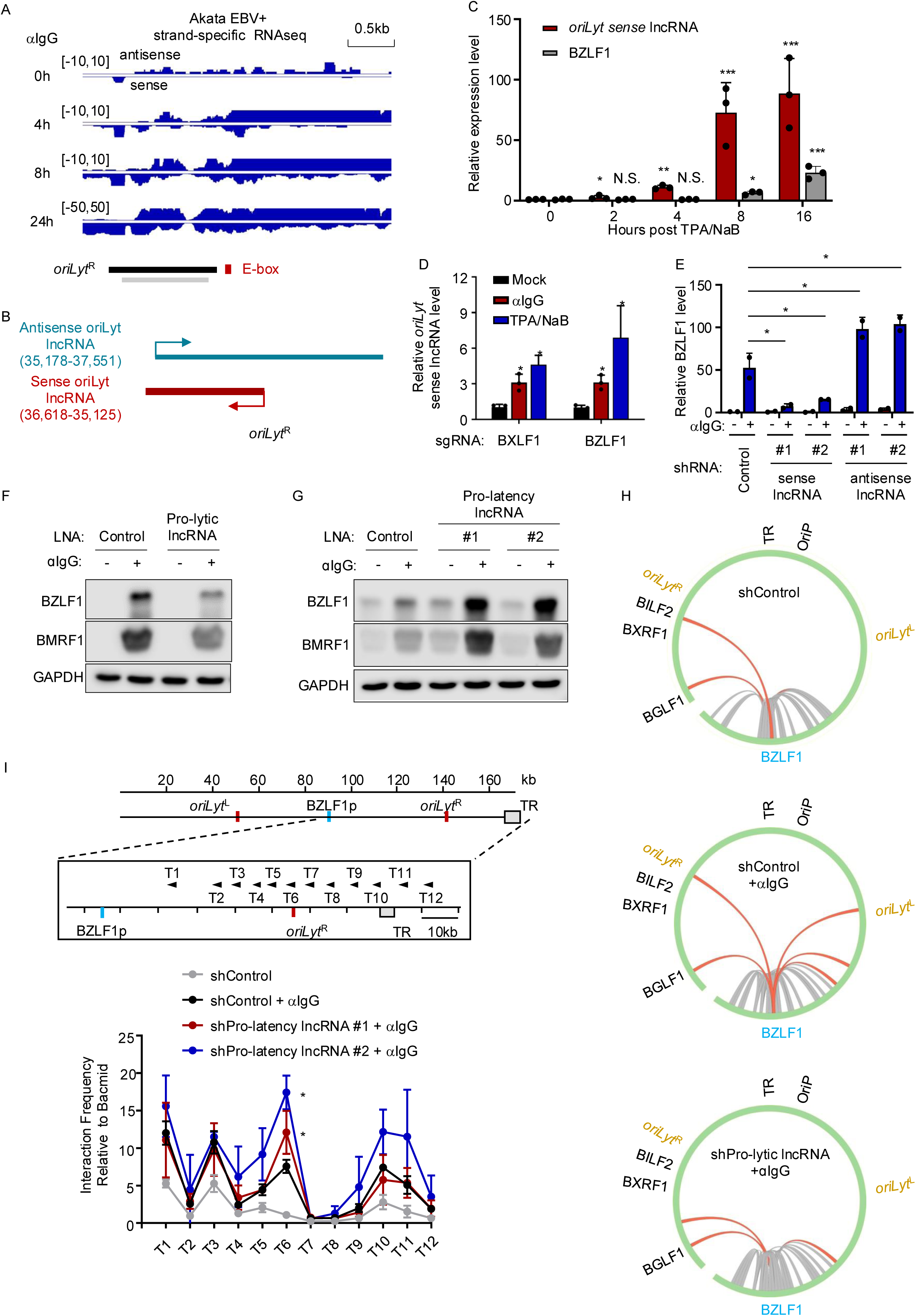
OriLyt lncRNA roles in EBV latency versus reactivation. (A) Strand-specific analysis^56^ of *oriLyt^R^* region transcripts in Akata cells stimulated by αIgG crosslinking for the indicated number of hours. Black bar depicts the *oriLyt^R^* region, grey bar indicates highly conserved *oriLyt^R^* and *oriLyt^L^* sequence. The *oriLyt* E-box highly occupied by MYC is indicated. (B) Schematic of oriLyt antisense and sense lncRNAs, as determined by 5’-and 3’- RACE analysis of Akata cells treated with TPA/NaB for 24h. (C) Mean ± S.D. values from 3 replicates of strand-specific RT-qPCR analysis of the oriLyt sense lncRNA and BZLF1 mRNA levels from Akata cells treated with TPA/NaB for the indicated hours. (D) Mean ± S.D. values from 3 replicates of strand-specific RT-qPCR analysis of oriLyt sense lncRNA level in Cas9+ Akata cells expressing sgRNAs targeting control *BXLF1* or *BZLF1*, mock stimulated or stimulated by αIgG or TPA/NaB for 4h. (E) Mean ± S.D. BZLF1 mRNA levels from values from 3 replicates of Akata cells with shRNAs targeting *oriLyt* sense or antisense lncRNAs and stimulated by αIgG for 24 hours. (F) Immunoblot analysis of WCL from Akata cells expressing control or *oriLyt* sense lncRNA targeting locked nucleic acid (LNA) and stimulated by αIgG for 24 hours. (G) Immunoblot analysis of WCL from Akata cells expressing control or *oriLyt* antisense lncRNA targeting LNA and stimulated by αIgG crosslinking for 24h. (H) Effects of oriLyt sense lncRNA knockdown on *BZLF1* promoter long-range interactions. H3K27ac HiChIP circos plots of long-range interactions between the *BZLF1* promoter and other EBV genome regions from Akata cells that expressed shRNA control or shRNA against the oriLyt pro-lytic lncRNA (bottom) and that were mock stimulated (top) or stimulated by αIgG for 24 hours. Red arcs indicate BZLF1 long-range (>20kb) interactions, gray and orange arcs indicate interactions <20kb. n=2 replicates. (I) Pro-Latency lncRNA knockdown effects on *BZLF1* promoter looping. Mean ± S.D. values from 3 replicates of 3C assays in Akata cells that expressed control shRNA or shRNA targeting the *oriLyt* pro-latency lncRNA, using a *BZLF1p* anchor and the indicated test primers. Significance differences refer to comparisons between Akata cells expressing control shRNA (black line) and shRNA targeting the *oriLyt* pro-latency lncRNA (red and blue lines) upon stimulation with αIgG. Blots are representative of 3 replicates. *, P<0.05

To determine *oriLyt* RNA transcription start sites (TSS) and termination sites, we performed 5’ and 3’-RACE (RNA amplification of cDNA ends). This identified the sense strand transcript induced upon reactivation as ∼1.4kb, originating from one of two TSS 52 bp apart (**Figure 3B** and **S3D**). The antisense strand transcript expressed in latency was found to be 2374 bases and to share the EBV LF3 promoter (**Figure 3B** and **S3D**). Two additional minor transcripts with TSS 1,139 bp downstream and with the same termination site were also identified. Next, we quantified the abundance of *oriLyt* transcripts in Akata EBV+ cells. During latency, each cell contained ∼160 copies of antisense *oriLyt* lncRNA, while 10-fold less *oriLyt* sense lncRNA was present (**Figure S3E-F**).

We used strand-specific RT-qPCR to define αIgG crosslinking effects on *oriLyt* lncRNA levels. In latent Akata cells, *oriLyt* sense but not antisense lncRNA level increased by 1 hour post αIgG treatment (**Figure S3G**). In Mutu Burkitt cells, sense *oriLyt* message was increased by 2 hours of TPA/NaB treatment, prior to upregulation of BZLF1 mRNA, and continued to increase through 16 hours post-treatment (**Figure 3C**). The antisense *oriLyt* lncRNA increased to a lesser extent by 8 hours post-TPA/NaB treatment (**Figure S3H**). Further suggestive of sense lncRNA induction preceding BZLF1 immediate early gene expression, we observed similar levels of the sense lncRNA message in Cas9+ Akata cells expressing sgRNAs targeting the BXLF1 and BZLF1 genes and then stimulated by αIgG or TPA/NaB (**Figure 3D**).

To determine if sense and antisense *oriLyt* lncRNAs were important for control of the EBV lytic switch, we expressed control or lncRNA targeting shRNA. Strand-specific RT-PCR confirmed successful on-target knockdown of either lncRNA (**Figure S3I**). Interestingly, sense lncRNA knockdown strongly impaired BZLF1 and BMRF1 induction by αIgG crosslinking, whereas antisense lncRNA knockdown had the opposite effect (**Figure 3E, S3J-K**). We therefore refer hereafter to the sense and antisense message as *oriLyt* pro-lytic and pro-latency lncRNAs, respectively. To further validate these findings, we identified two locked nucleic acids (LNA) that knocked down the pro-latency *oriLyt* lncRNA and one that depleted the pro-lytic *oriLyt* lncRNA. We again observed that pro-lytic lncRNA diminished EBV lytic reactivation by αIgG crosslinking, whereas pro-latency lncRNA knockdown had the opposite effect (**Figure 3F-G and S3L**).

We further explored effects of *oriLyt* lncRNA knockdown on EBV genomic structure. H3K27ac HiChIP highlighted that shRNA pro-Lytic (sense) *oriLyt* lncRNA knockdown significantly reduced *oriLyt/BZLF1* promoter looping triggered by αIgG crosslinking (**Figure 3H**). 3C analysis likewise revealed that pro-Lytic *oriLyt* lncRNA depletion significantly reduced *oriLyt:BZLF1* promoter contact frequency following αIgG crosslinking (**Figure 3I**), whereas pro-latency *oriLyt* lncRNA depletion had the opposite effect (**Figure S3M**). Collectively, these data support a model in which *oriLyt* encodes lncRNAs with opposing effects on the EBV lytic switch.

### MYC and the LSD1/CoREST/ZNF217 complex repress pro-Lytic lncRNA expression

We next characterized epigenetic control of pro-lytic lncRNA expression. CRISPR analyses identified MYC and the LSD1/CoREST/ZNF217 complex as key host factors that bind to *oriLyt* and suppress its looping to BZLF1^15,40^. Interestingly, MYC-occupied E-box sites are located adjacent to the pro-lytic *oriLyt* lncRNA TSS. We therefore tested MYC effects on *oriLyt* transcription, given also that MYC levels rapidly decrease upon EBV lytic reactivation^40,58^. MYC overexpression impaired pro-lytic *oriLyt* lncRNA induction by αIgG crosslinking, whereas CRISPR MYC KO was sufficient to upregulate pro-lytic lncRNA expression (**Figure 4A-B**). Consistent with this observation, enforced MYC expression also reduced pro-Lytic lncRNA promoter levels of total and serine 2 phosphorylated RNA Pol II at two hours post-αIgG treatment (**Figure S4A**). Taken together, these results suggest that MYC represses pro-lytic *oriLyt* lncRNA expression.

**Figure 4.**
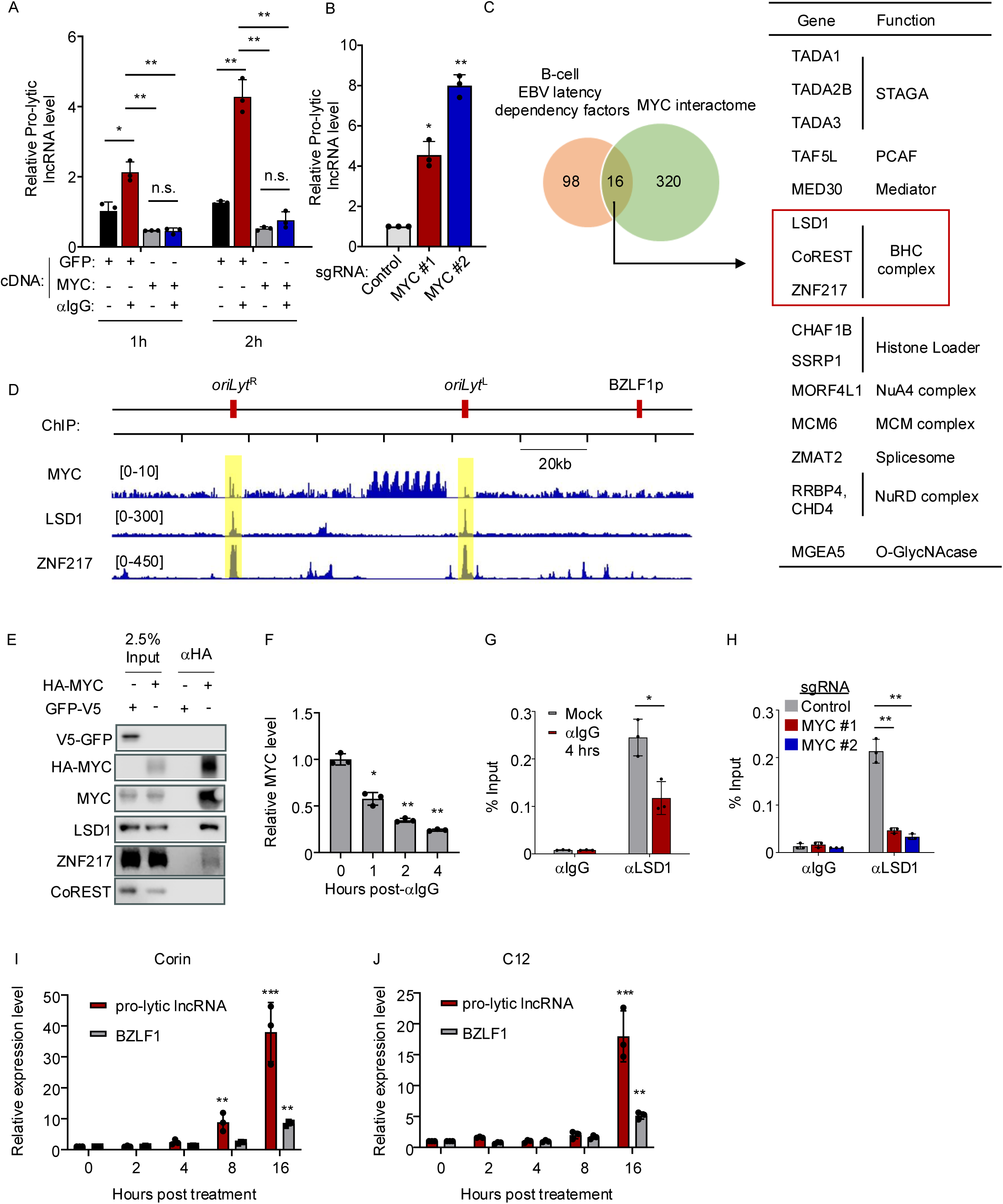
MYC supports LSD1/ZNF217/CoREST complex recruitment to *oriLyt* to repress pro-lytic *oriLyt* lncRNA expression. (A) Analysis of MYC overexpression blocks *oriLyt* pro-lytic lncRNA induction by αIgG. Mean ± S.D. *oriLyt* pro-lytic lncRNA levels from n=3 strand specific RT-PCR replicates using Akata cells that expressed GFP or HA-MYC encoding cDNAs and following mock or αIgG crosslinking for one or two hours. (B) MYC depletion upregulates oriLyt sense lncRNAs. Mean ± S.D. *oriLyt* pro-lytic lncRNA levels from n=3 RT-PCR replicates of Cas9+ Akata cells expressing control or MYC targeting sgRNA. (C) Cross-comparison of CRISPR-defined host factors necessary for EBV B cell latency^119^ and a BioID proximity labeling MYC interactome^120^. 16 hits common to both approaches are highlighted at right and included LSD1, CoREST and ZNF217. (D) ChIP-seq analysis of EBV genomic occupancy of MYC in EBV+ Daudi Burkitt cells^14^ versus LSD1 and ZNF217 in EBV+ Akata cells. Track heights are indicated at top. Data are average of 2 replicates. *oriLyt* regions are highlighted in yellow. (E) Immunoblot analysis of 2.5% input versus αHA immunoprecipitated complexes from Akata cells that expressed control HA-tagged GFP or MYC. Representative of n=3 replicates. (F) αIgG effects on EBV+ Burkitt MYC abundance. Mean ± S.D. GAPDH-normalized MYC mRNA levels from 3 RT-PCR replicates of Akata cells treated with αIgG for the indicated hours. (G) Effects of αIgG crosslinking on LSD1 *oriLyt* occupancy. Mean ± S.D. values from 3 ChIP-qPCR replicates of BZLF1 knockout Akata cells^18^, mock stimulated or stimulated by αIgG crosslinking for 4 hours, and using control IgG or αLSD1 antibodies and *oriLyt* primers. (H) Effects of MYC depletion on *oriLyt* LSD1 occupancy. Mean ± S.D. values from 3 ChIP-qPCR replicates, as in (G), using Akata cells expressing control or independent *MYC* targeting sgRNAs. (I) Effects of LSD1/HDAC inhibition on *oriLyt* pro-lytic lncRNA and BZLF1 expression. Mean ± S.D. pro-lytic lncRNA and BZLF1 mRNA levels from n=3 replicates of strand-specific qPCR on Akata cells treated with the dual LSD1/HDAC inhibitor corin (5 μM) for the indicated time. (J) Effects of LSD1 inhibition on *oriLyt* pro-lytic lncRNA and BZLF1 expression. Mean ± S.D. pro-lytic lncRNA and BZLF1 mRNA levels from 3 replicates of Akata cells treated with the LSD1 inhibitor C12 (5 μM) for the indicated time, as in (I).

While MYC:MIZ-1 complexes can repress target gene expression^59,60^, CRISPR screens have not implicated MIZ-1 in maintenance of EBV latency^15,40^. Indeed, CRISPR MIZ-1 KO did not derepress BZLF1 (**Figure S4B**), suggesting that alternative MYC interactors instead repress *oriLyt* pro-lytic lncRNA expression, together with MYC. We therefore cross-compared proximity labeling defined high confidence MYC interactors^61^ with our CRISPR hits, which identified 16 high-confidence MYC interactors critical for maintenance of EBV B cell latency (**Figure 4C**). Of these, we recently found that the LSD1/CoREST/ZNF217 complex occupies *oriLyt* in latency and prevented its looping to the *BZLF1* promoter^15^. Indeed, ChIP-seq analyses highlighted that ZNF217, LSD1 and MYC co-occupied *oriLyt* sites in latently EBV infected Burkitt cells (**Figure 4D**). Furthermore, CoREST and ZNF217 co-immunoprecipitated with MYC from Akata cell extracts (**Figure 4E**).

Upon IgG crosslinking or TPA/NaB treatment, MYC expression rapidly decreased (**Figure 4F** and **S4C-D**). Although EBV reactivation does not significantly change overall LSD1, CoREST or ZNF217 abundances^58^ (**Figure S4C**), *oriLyt* LSD1 occupancy rapidly decreased upon αIgG crosslinking, even in BZLF1 knockout Akata cells^18^ (**Figure 4G**). Furthermore, MYC knockout strongly diminished LSD1 *oriLyt* occupancy, indicating that MYC supports LSD1 recruitment to *oriLyt* (**Figure 4H**). In addition, CRISPR editing of MYC-bound E-box *oriLyt* sites, or depletion of MYC by CBL0137^40^, also diminished *oriLyt* LSD1 occupancy (**Figure S4E-F**). CRISPR KO of LSD1, CoREST or ZNF217 de-repressed EBV lytic proteins as we previously observed, but also induced pro-lytic *oriLyt* lncRNA expression (**Figure S4G-H**), as did LSD1 inhibition by the dual LSD1/HDAC inhibitor corin^62^ or the LSD1 inhibitor C12^63^ (**Figure 4I-J**). Taken together, these findings indicate that MYC and the LSD1/CoREST/ZNF217 complex together repress pro-lytic *oriLyt* lncRNA expression.

### Pro-lytic *oriLyt* transcription promotes CTCF binding and *oriLyt:BZLF1* looping

To test whether pro-lytic *oriLyt* lncRNA expression in *cis* drives reactivation, we used CRISPR-activation (CRISPRa), in which a sgRNA directs a CRISPR-dCas9 fused to a VP64 transcription activator (**Figure 5A**). CRISPRa successfully induced *oriLyt* pro-lytic lncRNA, BZLF1 and BMRF1 expression in Akata and MUTU I B cells (**Figure 5B-C, S5A-C**) and increased intracellular and encapsidated extracellular lytic EBV genome copy number, suggestive of a productive EBV lytic cycle (**Figure 5D**).

**Figure 5.**
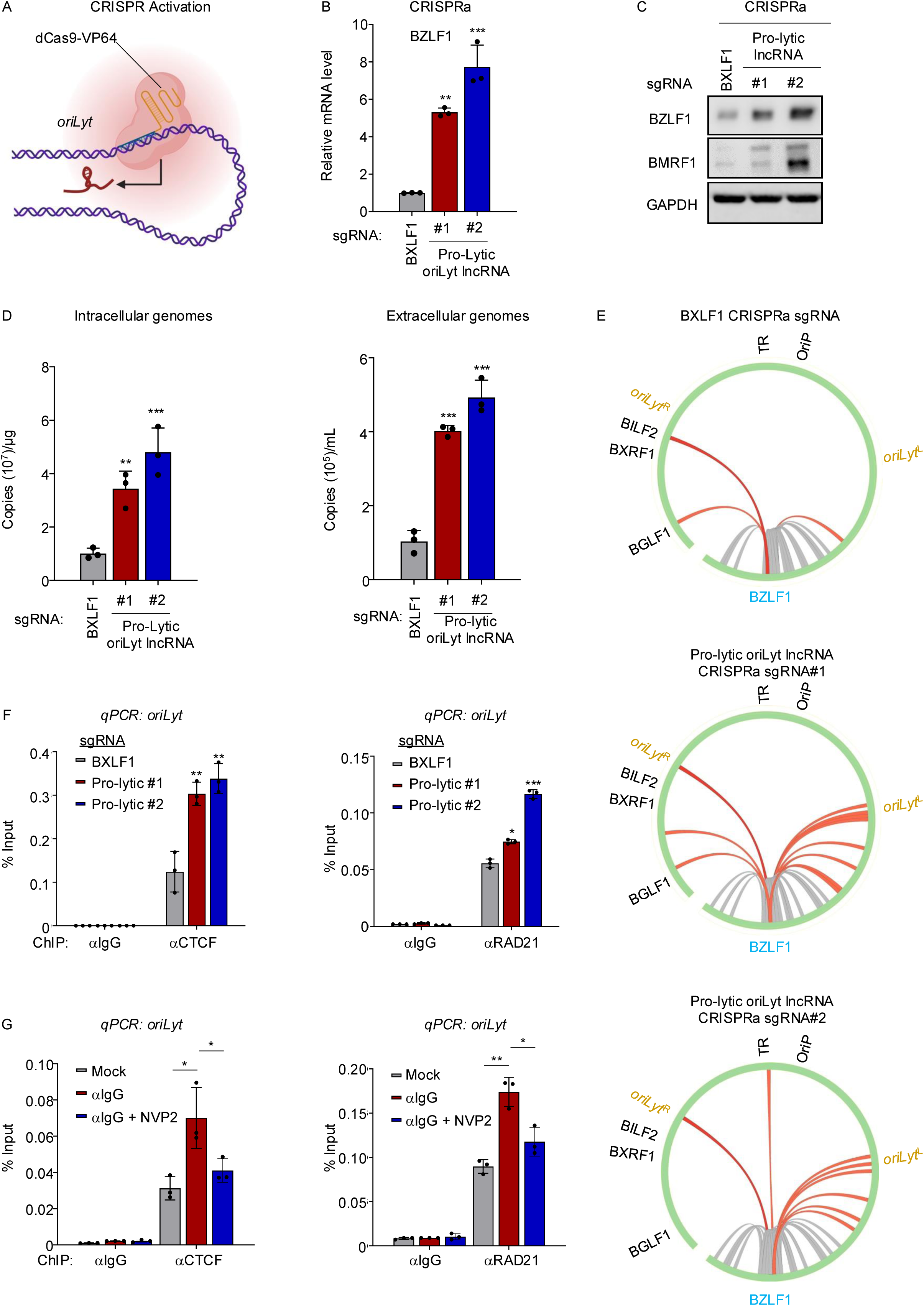
*OriLyt* transcription drives its looping to *BZLF1* and EBV reactivation. (A) Schematic model of CRISPR activation (CRISPRa) upregulation of *oriLyt* enhancer regions. sgRNA programs the dCas9/VP64 transcription activator. (B) CRISPRa driven *oriLyt* pro-lytic lncRNA expression induces BZLF1 mRNA expression. Mean ± S.D. GAPDH-normalized BZLF1 levels from 3 RT-PCR replicates of dCas9-VP64+ Akata cells with control BXLF1 vs *oriLyt* pro-lytic lncRNA promoter targeting sgRNAs. (C) *OriLyt* pro-lytic lncRNA expression induces BZLF1 protein expression. Immunoblot of WCL from Akata cells as in (B). Blots are representative of n=3 replicates. (D) O*riLyt* pro-lytic lncRNA expression induces EBV genome replication. Mean ± S.D. values from RT-qPCR analysis of EBV intracellular (left) or DNase-treated extracellular viral genome (right) copy number from Akata cells as in (B). (E) O*riLyt* pro-lytic lncRNA expression drives *oriLyt:BZLF1* promoter looping. Circos plots of H3K27ac HiChIP defined interactions between the *BZLF1* promoter and other EBV genome regions (orange arcs) in Akata cells as in (B) from n=2 replicates. (F) O*riLyt* pro-lytic lncRNA expression upregulates *oriLyt* CTCF occupancy. Mean ± S.D. values from ChIP-qPCR analysis of CTCF (left) and RAD21 (right) *oriLyt* occupancy in Akata cells as in (B). (G) αIgG crosslinking upregulates CTCF and RAD21 *oriLyt* occupancy in a transcription-dependent manner. Mean ± S.D. values from ChIP-qPCR analysis of CTCF (left) and RAD21 (right) *oriLyt* occupancy in Akata cells mock stimulated or stimulated by αIgG for 4 hours ± the Pol II antagonist NVP2 (0.5nM). *P<0.05, **P<0.01.

We similarly tested pro-latency *oriLyt* lncRNA upregulation effects on EBV lytic gene expression. In contrast to effects of pro-lytic lncRNA induction, CRISPRa pro-latency lncRNA upregulation did not induce lytic gene expression and diminished BZLF1 induction by αIgG crosslinking (**Figure S5D**). These results suggest that the pro-latency lncRNA can fine-tune lytic gene levels. We next asked if CRISPRa pro-lytic lncRNA upregulation was sufficient to drive *oriLyt:BZLF1* promoter DNA looping. We performed H3K27ac HiChIP following CRISPR activation by control *BXLF1* versus either of two pro-lytic *oriLyt* lncRNA targeting sgRNAs. Interestingly, CRISPRa pro-lytic lncRNA upregulation driven by either of two sgRNAs caused long-range interactions between the *BZLF1* promoter region and both *oriLyt* regions (**Figure 5E**), which were also detected by 3C assays (**Figure S5E**).

Given its effects on DNA looping, we hypothesized that the pro-lytic lncRNA alters CTCF and/or cohesin occupancy at *oriLyt*. To test this, we measured effects of CRISPRa mediated pro-lytic lncRNA upregulation on CTCF and RAD21 cohesin occupancy at *oriLyt*. CRISPR activation of pro-lytic lncRNA expression by either of two sgRNAs significantly increased CTCF and RAD21 occupancy at *oriLyt* (**Figure 5F**). To therefore test if transcription was necessary for increased CTCF and RAD21 deposition at *oriLyt* upon B cell receptor crosslinking, we treated cells with the CDK9 inhibitor NVP2^64^ to block Pol II elongation. As expected, NVP2 blocked αIgG crosslinking upregulation of either the pro-lytic oriLyt lncRNA or BZFL1 (**Figure S5F-G**). Interestingly, NVP2 almost completely disrupted αIgG crosslinking-driven CTCF and RAD21 loading onto *oriLyt* and also *oriLyt:BZLF1* promoter DNA looping (**Figure 5G** and **S5H**). Interestingly, NVP2 also decreased BZLF1 promoter RAD21 but not CTCF occupancy (**Figure S5H**). These results suggest that the oriLyt pro-lytic lncRNA interacts with DNA looping machinery to support rapid formation of a DNA loop that juxtaposes an EBV genomic enhancer with the immediate early gene promoter.

### The pro-lytic *oriLyt* lncRNA contains an RNA G-quadruplex that recruits CTCF

To further investigate how *oriLyt* lncRNAs reciprocally control the viral B cell lytic switch, we performed chromatin isolation by RNA purification (ChIRP) sequencing^65^, which maps genome-wide lncRNA binding sites. ChIRP-seq revealed that both lncRNAs occupy *oriLyt*^R^ and *oriLyt* ^L^ regions (**Figure 6A**), but not other EBV or host genomic sites. Interestingly, within the ChIRP-seq limits of detection, the pro-lytic lncRNA occupied the *oriLyt* CBS to a greater degree than the pro-latency lncRNA, particularly at oriLyt^L^ (**Figure S6A**). Both *oriLyt* lncRNAs, LSD1 and ZNF217 highly co-occupied *oriLyt* sites, suggesting possible cross-talk between them (**Figure S6A**). To next investigate whether the *oriLyt* lncRNA could also work when over-expressed exogenously. However, lentiviral U6 promoter driven pro-lytic lncRNA over-expression only mildly induced EBV lytic gene expression (**Figure S6B-C**).

**Figure 6.**
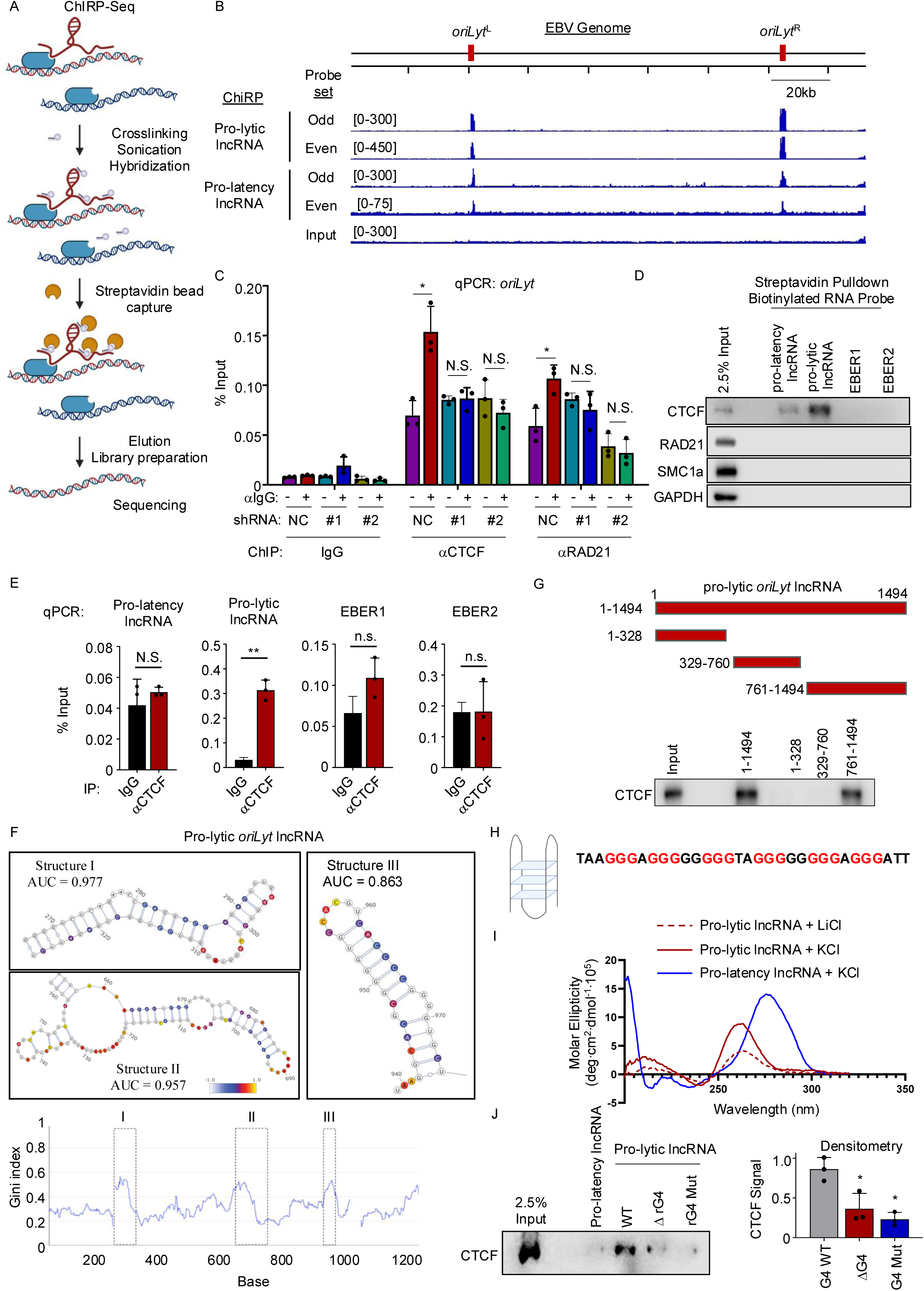
A pro-lytic lncRNA encoded RNA G4 quadruplex recruits CTCF to oriLyt. (A) ChIRP-seq schematic diagram. (B) ChIRP-seq analysis of *oriLyt* lncRNA EBV genomic occupancy following αIgG crosslinking for 24h. Average of 2 replicates. (C) oriLyt pro-lytic lncRNA knockdown reduces *BZLF1* promoter CTCF and RAD21 occupancy. Mean ± S.D. values from 3 ChIP-qPCR replicates of CTCF and RAD21 *oriLyt* occupancy. Akata cells expressed shRNA non-targeting control (NC) or shRNA targeting the *oriLyt* pro-lytic lncRNA and were treated with αIgG for 4hours as indicated. (D) CTCF associates with the pro-lytic lncRNA. 25pmol biotin-labeled *oriLyt* pro-latency lncRNA, pro-lytic lncRNA, EBER1 or EBER2 were co-incubated with Akata cell lysates for 4 hours. Input vs streptavidin purified complexes were analyzed by immunoblot. (E) Pro-lytic lncRNA CTCF association. Mean ± S.D. strand-specific RT-PCR values from 3 replicates of immunoglobulin control vs CTCF complexes immunopurified from nuclear extracts of αIgG stimulated Akata cells. (F) DMS-MaPseq pro-lytic lncRNA structural analysis. Pro-lytic lncRNA GINI index scores, where higher numbers indicate structured regions. Several predicted lncRNA structures are shown above with their corresponding Area Under the Receiver Operating Curve (AUC). AUC is a measure of agreement between the DMS reactivity data and the predicted structure model; a value of 1.0 indicates perfect agreement. (G) A pro-lytic lncRNA 3’ region mediates CTCF association. Immunoblot analysis as in (D) of streptavidin purified biotinylated full length or truncated pro-lytic lncRNA complexes. (H) Pro-lytic lncRNA predicted RNA G-quadruplex (rG4) sequence. (I) Circular dichroism (CD) analysis of *in vitro* transcribed lncRNA. CD spectra of *oriLyt* pro-latency or pro-lytic lncRNAs in 150mM KCl buffer or of the pro-lytic lncRNA in 150mM LiCl buffer. The pro-lytic lncRNA positive peak at 260nm and negative peak at 240nm in KCl buffer which decreased in LiCl buffer are indicative of an rG4 structure. Representative of 3 replicates. (J) The pro-lytic lncRNA rG4 is critical for CTCF association. CTCF immunoblot analysis of streptavidin purified biotinylated lncRNA complexes from Akata extracts, as in (G). βG4, pro-lytic lncRNA G4 deletion mutant; rG4 Mut, rG4 G mutated to non-G nucleotides. Densitometry summarizes CTCF signals from 3 replicates. Blots in D, G and J are representative 3 independent replicates. *P<0.05, n.s, non-significant.

Enhancer RNAs can physically interact with the CREB-binding protein (CBP) to activate its histone acetyltransferase activity^66^. This interaction potentially connects eRNA transcription with H3K27 acetylation during enhancer activation^67^. Thus, we tested if the pro-lytic lncRNA regulates EBV genome structure by modulating epigenetic states. However, pro-lytic lncRNA knockdown did not significantly alter H3K27ac or H3K4me1 levels in αIgG crosslinked Akata cells (**Figure S6D**). Rather, pro-lytic lncRNA knockdown diminished *oriLyt* CTCF and RAD21 occupancy following αIgG crosslinking (**Figure 6C**). Therefore, the pro-lytic lncRNA acts at the level of DNA looping factor recruitment to *oriLyt*, rather than enhancing *oriLyt* enhancer activity.

CTCF zinc fingers bind to RNA, and this association is required for CTCF-mediated chromatin loop formation^68^. Furthermore, we noticed that *oriLyt* CBS have lower DNA sequency similarity to the consensus CBS motif than the *BZLF1* promoter CBS, suggesting that the former may be a low-affinity site that interacts more weakly with CTCF. These observations prompted us to investigate if the pro-lytic lncRNA facilitates CTCF binding to the *oriLyt* CBS. We incubated *in vitro* transcribed and biotinylated pro-lytic lncRNA, pro-latency lncRNA, or as negative controls EBV-encoded EBER1 or EBER2 non-coding RNAs with Akata nuclear extracts. Immunoblot analysis of streptavidin purified complexes highlighted that the pro-lytic lncRNA, but not the other EBV RNAs, associated with CTCF (**Figure 6D**). By contrast, it did not co-purify RAD21 or SMC1a cohesins or GAPDH, suggesting specificity of the pro-lytic lncRNA/CTCF complex. Similarly, we tested whether we could detect EBV RNAs associated with endogenous CTCF. The pro-lytic lncRNA, but not the pro-latency lncRNA or EBER RNAs, co-immunopurified from αIgG crosslinked Akata cell extracts, despite each of their abundant expression in Akata cells (**Figure 6E**).

We then explored which pro-lytic lncRNA region associated with CTCF. We probed the pro-lytic lncRNA structure in Burkitt cells using Dimethyl sulfate (DMS). DMS MaPseq analysis^69^ revealed 3 highly structured lncRNA regions, based on high Gini index scores (**Figure 6F**). To determine which region of the lncRNA interacts with CTCF, we designed pro-lytic lncRNA deletion mutants and performed biotinylated RNA pulldown analysis with nuclear extracts from αIgG crosslinked Burkitt cells. This highlighted that lncRNA bases 761-1494, but not bases 1-328 or 329-760, associated with CTCF to a similar degree as the full length lncRNA (**Figure 6G**).

We noticed that the pro-lytic lncRNA contains a guanine-rich region with putative RNA G-quadruplex (rG4) forming sequence (GGGN1-7GGGN1-7GGGN1-7GGG, where N is any base) between bases 960-1146, within the lncRNA fragment that retained CTCF association (**Figure 6H**). Guanine-rich RNA sequences with consecutive guanines can form quadruple-stranded rG4 structures, which are stabilized by monovalent cations, such as potassium. Importantly, multiple studies highlight that CTCF binds particularly strongly to rG4 structures, which for instance play major roles in Xist lncRNA mediated CTCF recruitment for X-chromosome silencing^70,71^. Therefore, we explored the possibility that the pro-lytic lncRNA uses an rG4 structure to recruit CTCF to *oriLyt*. Circular dichroism (CD) spectroscopy analysis of *in vitro* transcribed pro-lytic lncRNA revealed a positive peak at ∼260nm and a negative peak at ∼240nm (**Figure 6I**), characteristic of a parallel rG4 structure. As a positive control, we observed similar CD signals with the telomeric TERRA lncRNA (**Figure S6E**)^72,73^, but not with the pro-latency oriLyt lncRNA (**Figure 6I**). Moreover, the 240nm peak decreased when the pro-lytic lncRNA was incubated in buffer with lithium (Li+) rather than potassium (K+), further supporting the presence of a pro-lytic lncRNA rG4 structure (**Figure 6I, S6E**). The pro-lytic lncRNA rG4 was also detectable *in vivo* following EBV reactivation by an RNA immunoprecipitation-qPCR approach: rG4 immunoprecipitation with the antibody BG4^74^ followed by qPCR revealed enrichment for the pro-lytic RNA, but not for negative control EBER2 following Akata B cell αIgG crosslinking or TPA/NaB treatment (**Figure S6F**).

To define whether the pro-lytic lncRNA rG4 was necessary for its association with CTCF, we co-incubated biotinylated wildtype, rG4 deletion mutant or rG4 point mutant (where ten guanines were mutated to other bases) pro-lytic lncRNA with Akata nuclear extracts. Streptavidin pulldown analysis revealed that wildtype, but not rG4 mutant pro-lytic lncRNA, associated with CTCF (**Figure 6J**). We therefore investigated whether the rG4 structure was also important for lytic gene expression and CTCF association *in vivo*. The cationic porphyrin TMPyP4, which unfolds rG4 and disrupts rG4-dependent protein interactions^75,76^ disrupted pro-lytic lncRNA rG4 structure in a dose-dependent manner (**Figure S7A**) and inhibited EBV lytic gene expression induced by IgG-crosslinking (**Figure S7B**). TmPyP4 also impaired pro-lytic lncRNA association with CTCF and downmodulated *oriLyt:BZLF1* promoter interaction in αIgG-crosslinked Akata cells (**Figure S7C-D**). Collectively, these results indicate that rapidly induced pro-lytic lncRNA rG4 recruits CTCF to *oriLyt*, which then drives rapid higher order EBV genomic reorganization to induce reactivation.

### Antagonism between the pro-latency *and* pro-lytic oriLyt lncRNA

We next explored how the pro-latency *oriLyt* lncRNA mechanistically counteracts EBV reactivation. Building on our observation that its overexpression in *cis* impairs EBV reactivation, we tested the hypothesis that it impairs CTCF loading at *oriLyt*. Indeed, pro-latency *oriLyt* lncRNA knockdown increased CTCF occupancy at both *oriLyt* and at the *BZLF1* promoter, both in latency and following αIgG-crosslinking (**Figure 7A**). We therefore suspected that the pro-latency lncRNA may function by forming a duplex with the pro-lytic RNA to serve as a sponge and potentially also to prevent rG4 formation to mask its association with CTCF. To survey for duplex formation *in vivo*, we used psoralen crosslinking followed by nuclease digestion and proximity RNA ligation^77,78^ on αIgG-crosslinked Burkitt cells. The presence of ligase-induced artificial junctions suggested the presence of *in vivo* duplex formation, which were significantly reduced by LNA-mediated antisense oriLyt lncRNA knockdown, suggestive of specificity (**Figure S7E**).

**Figure 7.**
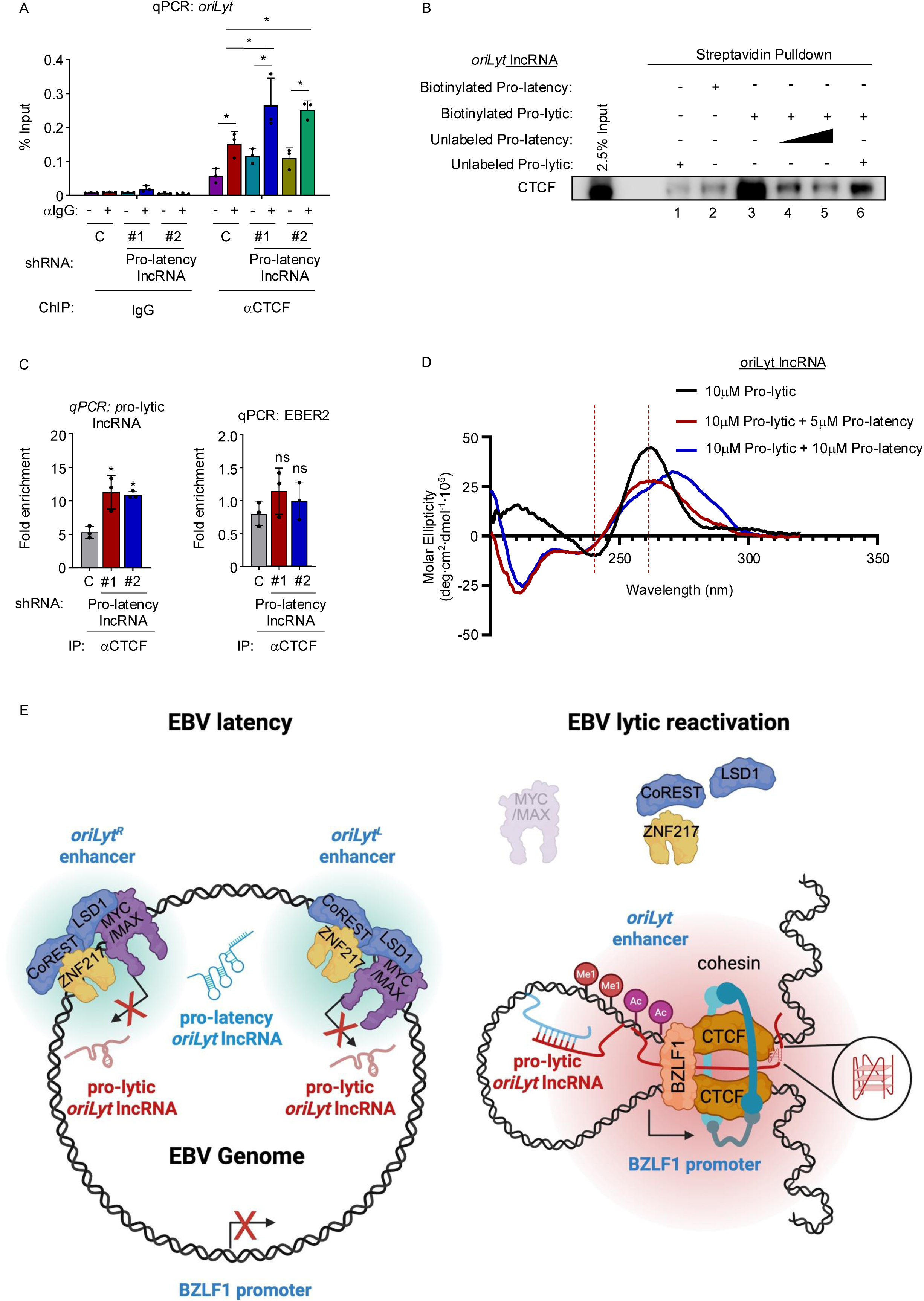
The pro-latency *oriLyt* lncRNA negatively regulates pro-lytic lncRNA rG4 assembly and CTCF recruitment. (A) Pro-latency lncRNA knockdown increases *oriLyt* CTCF occupancy. Mean CTCF and pro-lytic lncRNA (B) The pro-latency lncRNA impedes pro-lytic lncRNA association with CTCF. Biotinylated lncRNAs (25pmol), alone or together with unbiotinylated lncRNAs (12.5 pmol or 25pmol) and mixed with Akata cell extracts. Streptavidin purified complexes were analyzed by CTCF immunoblot. Representative of 3 replicates. (C) Pro-latency lncRNA knockdown increases CTCF and pro-lytic lncRNA association. Mean ± S.D. values from 3 replicates of fold enrichment of αCTCF RIP over IgG control RIP in strand-specific RT-qPCR for pro-lytic lncRNA (left) or EBER2 (right), using Akata cells that expressed the indicated shRNAs and that were stimulated by aIgG crosslinking for 4 hours. The RT-qPCR value derived from IgG control RIP was set to fold enrichment of 1. (D) Pro-latency lncRNA disrupts the pro-lytic lncRNA rG4. CD spectra of *oriLyt* pro-lytic and pro-latency lncRNA at the indicated ratios in 10mM LiCac buffer (pH 7.2) with 150mM KCl. (E) oriLyt lncRNA control of the B cell lytic switch. In latency, the pro-latency lncRNA is expressed, while MYC:MAX and LSD1/CoREST/ZNF217 repress pro-lytic lncRNA expression. Reactivation stimuli perturb transcription factor occupancy, enabling rapid *oriLyt* H3K4 methylation and *oriLyt* pro-lytic lncRNA expression. The pro-lytic lncRNA uses an rG4 tor recruit CTCF to oriLyt, which promotes cohesin mediated looping to the BZLF1 promoter. oriLyt enhancer juxtaposition with the BZLF1 promoter triggers the lytic cycle. *p<0.05.

We next asked if the pro-latency lncRNA competitively inhibited association between CTCF and the pro-lytic lncRNA. In support, pre-incubation with unlabeled pro-latency *oriLyt* lncRNA significantly decreased the level of CTCF that co-purified from Burkitt extracts with biotinylated pro-latency *oriLyt* lncRNA (**Figure 7B**). We then performed RNA immunoprecipitation assays to test effects of pro-latency oriLyt lncRNA knockdown on pro-lytic lncRNA/CTCF association in Burkitt cells. Pro-latency lncRNA knockdown significantly increased levels of pro-lytic lncRNA that co-immunopurified with CTCF, but not of EBER2 (**Figure 7C**). Furthermore, CD spectroscopy revealed that co-incubation with increasing amounts of pro-latency lncRNA reduced pro-lytic lncRNA rG4 formation, as evidenced by reduced 260nm positive and 240 nm negative signals (**Figure 7D**). Instead, co-incubation with the pro-latency lncRNA caused negative peak formation at 210nm, indicative of RNA duplex formation. Collectively, these results support a model in which the antisense *oriLyt* lncRNA is expressed in latency and sponges leaky sense *oriLyt* lncRNA to prevent its rG4 formation and CTCF association. Upon lytic induction stimuli, MYC, LSD1, CoREST and ZNF217 occupancy rapidly diminish, licensing swift pro-lytic lncRNA upregulation. This serves to overcome pro-latency lncRNA suppression, drive CTCF loading at *oriLyt* and its association with the *BZLF1* promoter, and to thereby juxtapose an EBV genomic enhancer with the B cell immediate early gene promoter to trigger reactivation (**Figure 7E**). Once BZLF1 is induced, it binds to seven *oriLyt* sites to convert it into a strong enhancer, completing a rapid positive feedback loop.

## Discussion

A biphasic lifecycle, with alternation between states of latency and reactivation, is a herpesvirus’s hallmark that enables persistent host infection. EBV establishes tight latency in newly infected B-cells, and upon reaching the memory cell reservoir, can even shuts off all viral protein expression^79^. Therefore, EBV latency is maintained by epigenetic mechanisms, as are the earliest events in reactivation, which is necessary for spread, but which risks immune detection. Lytic reactivation is increasingly implicated as a driver of a surprising range of EBV-associated malignancies and autoimmune diseases. However, key molecular details of how the EBV lytic switch is rapidly activated have remained incompletely understood.

Here, we identify two *oriLyt* enhancer lncRNAs and define their opposing roles in B-cell lytic switch regulation. We present evidence that the pro-lytic lncRNA is rapidly induced by lytic induction stimuli, by MYC or LSD1 perturbation, and uses an RNA G4 quadruplex to recruit the DNA looping factor CTCF to *oriLyt* enhancers. Pro-lytic lncRNA expression drove cohesion-mediated and CTCF-gated DNA looping to juxtapose *oriLyt* enhancers and the *BZLF1* promoter. Degron-tagged RAD21 depletion suggests that this is a key transcription stimulus for reactivation. The pro-latency lncRNA also localized to *oriLyt* and negatively regulated pro-lytic rG4 assembly and CTCF association, thereby serving as a sponge to establish a reactivation threshold. In this manner, *oriLyt* lncRNAs serve as molecular signals that integrate B-cell state to govern the viral lytic switch.

*oriLyt* have well-defined roles in late gene expression and in viral lytic DNA replication^10^. However, our HiChIP and 3C analyses identified that both *oriLyt* enhancers rapidly loop to the *BZLF1* promoter region upon B-cell receptor activation. Since we also found that cohesin-mediated looping strongly supports EBV immediate early gene expression, and that CRISPR-inhibition of *oriLyt* blocked whereas CRISPR-activation drove EBV reactivation, our data reveal key *oriLyt* enhancer roles in EBV reactivation. Once produced, BZLF1 binds to seven *oriLyt* response elements^80^ to upregulate *oriLyt* enhancer strength. Thus, we anticipate that the pro-lytic lncRNA triggered DNA loop drives an initial wave of BZLF1, which then strengthens the *oriLyt* enhancer and creates a positive feedback loop to further drive reactivation. Reliance upon viral lncRNAs to govern the lytic switch may enable latently infected cells to evade T-cell detection until immunomodulatory lytic cycle genes are expressed.

The ∼170 kb EBV genome approaches the size of eukaryotic genome topologically associated domains (TAD) genomic units that facilitate DNA looping interactions^81^. Furthermore, CBS are placed adjacent to both *oriLyt* and to the *BZLF1* promoter, which are distributed at roughly equidistant locations across the EBV genome, and RNA polymerase activity at CBS increases with EBV reactivation^57^. Therefore, we propose that EBV genomic architecture enables the pro-lytic lncRNA to control *oriLyt* CTCF loading to then rapidly remodel higher order EBV genomic structure by taking advantage of the remarkable speed of cohesin-mediated DNA looping. Since CTCF-bound EBV genomic loops also contribute to latency program selection^57,82-86^, we predict that distinct viral lncRNAs may also drive their formation. We further suggest that lncRNA mediated genome looping may be a theme that drives reactivation across herpesviruses.

EBV evolved complex epigenetic mechanisms to dictate pro-lytic lncRNA in response to environmental cues, such as B-cell receptor signaling. We present evidence that EBV evolved an unusual mechanism by which MYC and the LSD1/CoREST/ZNF217 complex repress pro-lytic lncRNA expression. MYC depletion, which occurs upon EBV+ Burkitt B-cell IgG crosslinking and also upon plasma cell differentiation. While primary human B-cell plasma cell differentiation models of EBV reactivation are not available, Burkitt MYC knockout or LSD1 inhibition each rapidly increased pro-lytic lncRNA expression, whereas MYC overexpression blocked its induction by αIgG crosslinking. An important future objective will therefore be to define how MYC and LSD1/CoREST/ZNF217 co-repress pro-lytic lncRNA expression, and whether the pro-latency lncRNA supports their action. Such a mechanism could enable assembly of a MYC-based repressor at *oriLyt*, but not at many host genomic MYC-bound sites.

ChIRP-seq highlighted that both the pro-latency and pro-lytic lncRNAs co-occupy both *oriLyt* regions. How they achieve this localization remains to be defined. They may bind to sequence specific host factors for recruitment to *oriLyt*, though their selectivity suggests that they may instead bind directly to *oriLyt* DNA in a sequence-specific fashion. Likewise, our data highlights that the pro-lytic lncRNA upregulates *oriLyt* CTCF occupancy. It will be of significant interest to determine by ChIRP and proteomic whether the pro-lytic lncRNA also recruits transcription factors or epigenetic enzymes to further promote immediate early gene transcription.

The pro-latency lncRNA establishes a barrier to reactivation by sponging the pro-lytic lncRNA and preventing its rG4 assembly. This may allow the pro-latency lncRNA to set a threshold level of pro-lytic lncRNA expression needed for reactivation in order to buffer against leaky pro-lytic lncRNA expression. This may be particularly important in newly infected cells as latency is established. Similarly, pro-latency lncRNA may have key latency gating roles on the single cell level, where MYC expression levels vary considerably, including across the cell cycle^87-89^ and even within EBV-transformed B cell populations^36^, to ensure that reactivation happens only beyond a threshold level of MYC depletion, as happens with plasma cell differentiation. STAT3 has also been reported to negatively regulate EBV reactivation^90-92^, and it will be of interest to determine its effects on EBV lncRNA expression.

How specific DNA anchor sites are selected for loop creation, and how anchors shift to form new loops in a physiologically relevant manner are not fully understood, particularly for EBV. The ‘CTCF Code’ hypothesizes that combinatorial usage of the 11 CTCF zinc fingers (ZnF) contributes to its binding and activity at specific sites^93,94^. CTCF is a G4-binding protein^70,95^, though most studies have focused on RNA/DNA hybrid G4 structure roles in CTCF recruitment to specific genomic sites. rG4 are more stable than DNA G4^96^, and CTCF may preferentially interact with rG4-containing RNA^70^. Therefore, we hypothesize that a key pro-lytic lncRNA feature is use of its rG4 as a bait to recruit and position CTCF at *oriLyt*.

EBV lytic reactivation is increasingly associated with both autoimmunity and cancer. EBV reactivation may be a key driver of multiple sclerosis and primary sclerosing cholangitis^7,97-99^. Similarly, EBV reactivation contributes to lymphomagenesis^34,100,101^ and may have tumor microenvironmental roles. Hyper-lytic EBV strains may contribute to carcinogenesis^102^. It will therefore be of interest to investigate pro-latency and pro-lytic oriLyt lncRNA expression in clinical samples across these contexts, and to test the hypothesis that changes in their expression may by associated with specific disease phenotypes.

Available chemotherapies do not differentiate between EBV+ and EBV- tumors and have significant morbidity. Few options are available for relapsed disease. Consequently, there is growing interest in modulating EBV reactivation for therapeutic applications. With regards to lymphoma and carcinoma, better approaches are needed to drive EBV reactivation in order to sensitize tumor cells to the highly cytotoxic effects of ganciclovir by the early lytic EBV kinase BGLF4^25^. It may be possible to harness pro-latency lncRNA knockdown, for example by lipid nanoparticle LNA delivery, to enhance reactivation responses to availble epigenetic modulators such as arginine butyrate^103^, nanatinostat^104^ or corin. Alternatively, it should be possible to harness pro-lytic lncRNA knockdown approaches to downmodulate EBV lytic gene expression in both the autoimmuity ad cancer settings.

In summary, we identified *oriLyt* encoded lncRNAs with opposing roles on the EBV lytic switch. Transcription of the pro-lytic lncRNA is repressed by key CRISPR-defined latency dependency factors. Once expressed, the pro-lytic lncRNA uses an rG4 to recruit CTCF to the *oriLyt* enhancer and to promote its looping to the BZLF1 immediate early promoter to drive reactivation. The pro-latency lncRNA serves as a sponge for the pro-lytic lncRNA, blocking assembly of its rG4 to establish a threshold level of pro-lytic lncRNA expression for reactivation. Targeting these lncRNAs may lay the foundation for EBV-targeted therapeutic approaches of EBV-associated malignancies and autoimmune diseases.

## Declaration of interests

The authors declare no competing interests.

## Ethics approval

Research with published EBV+ Burkitt and nasopharyngeal cell line models, whose origin is stated in the Star Methods section. This research was approved by our institutional review board and followed ethical guidelines like the Declaration of Helsinki.

## Data and code availability

HiChip, ChIRP-seq and ChIP-seq data will be deposited to the NIH Geo omnibus and will be released upon publication. No new code was written for this study.

## Acknowledgments

This work was supported by NIH R01AI164709, R01CA228700, U01CA275301, P01CA269043 to B.E.G. and by a Lymphoma Research Foundation Postdoctoral Fellowship to Y.L. We thank Karen Adelman and Hendrik Michel for helpful suggestions.

## Author contributions

Z.L., Y.L., S.L.G., G.R.A., and L.A.M.N. performed experiments and data analysis; W.D. and D.M. performed bioinformatic analysis; B.E.G., R.G., B.Z., S.R. and M.T. supervised the study. Z.L., Y.L. and B.E.G. wrote the manuscript.

## Star Methods

### Cell lines and cultures

The EBV+ Akata and Mutu I cell lines were obtained from Elliott Kieff and Jeff Sample and stably expressed *Streptococcus pyogenes* Cas9 or catalytically dead *Streptococcus pyogenes* Cas9 (dCas9) fused to repressor domain KRAB (dCas9-KRAB) for CRISPRi studies or dCas9 fused to the VP64 transactivation domain (dCas9-VP64) for CRISPRa studies. Briefly, *Streptococcus pyogenes* Cas9, dCas9-KRAB or dCas9-VP64 was introduced into Burkitt cell lines by lentivirus transduction followed by blasticidin selection (5μg/mL). Burkitt cells were grown in RPMI-1640 medium (Gibco, Life Technologies) supplemented with 10% fetal bovine serum (FBS, Gibco) and cultured at 37°C with 5% CO2 in a humidified chamber. MycoAlert Mycoplasma Detection Kit (Lonza) was routinely used to ensure that cell lines were mycoplasma free. HEK293T cells were obtained from ATCC and cultured in Dulbecco’s Modified Eagle Medium (DMEM) (Gibco) supplemented with 10% FBS.

### Plasmid generation

The RAD21 cDNA entry vector was purchased from DNASU. FKBP12 ^F36V^ was synthesized by IDT and fused to the C-terminal of RAD21 by HiFi DNA Assembly (NEB). The mutated PAM site of RAD21-FKBP12^F36V^ was introduced using Q5 Site-Directed Mutagenesis Kit (NEB) using primer set (RAD21 pam F/R, listed in **Supplementary Table 1**). RAD21-FKBP12^F36V^ with PAM site mutation was cloned into the Gateway destination vector pLX-TRC313 vector carrying V5 tag. To generate the *BZLF1* promoter firefly luciferase reporter vector, *BZLF1* promoter region bases -950 to +50 was PCR amplified and cloned into pGL3 (Promega). The *oriLyt* pro-latency lncRNA, full length and truncated *oriLyt* pro-lytic lncRNA, the rG4-deleted and the rG4-mutated *oriLyt* pro-lytic lncRNA were generated by PCR amplification and cloned into pcDNA3.1(-) for *in vitro* transcription or electroporation. Briefly, primers were designed to amplify 3 truncated regions (1-328nt, 329-760nt and 761-1494nt), full length with deleted rG4 motif (TAAGGGAGGGGGGGGTAGGGGGGGGAGGGATT) and full length with mutated rG4 motif (TAAGTGAGATCCGTGTAGATGGGTGAGTGATT). PCR fragments were fused into digested pcDNA3.1(-) vector using HiFi DNA Assembly (NEB). All primers were listed in the **Supplemental Table 1-3**.

### Construction of RAD21-FKBP12 ^F36V^ Akata cell line

RAD21-FKBP12^F36V^ Akata cells were generated by introducing pLX-TRC313-RAD21-FKBP12^F36V^ with PAM site mutation into Cas9+ Akata EBV+ cells followed by hygromycin selection for 14 days. To deplete endogenous RAD21, cells were transduced with lentivirus that expressed a RAD21 targeting sgRNA followed by puromycin selection (3 μg/mL) for 7 days. Single cell clones were selected and screened for biallelic endogenous RAD21 knockout.

### Electroporation

Locked nucleic acids (LNA) were designed and purchased from IDT. Culture media was pre-warmed in 37℃ for 0.5h. 5×10^5^ Akata EBV+ cells were washed with PBS and resuspended in 10 μl R buffer. 100pmol LNA or 1µg plasmid was electroporated by an Invitrogen Neon Transfection System (1350V, 30ms, 1 pulse). LNAs sequences are listed in Supplemental Table (**Supplemental Table 4**).

### HiChIP

HiChIP was done as previously published^105^. Briefly, 10 million Akata cells were harvested and crosslinked with freshly made 1% formaldehyde (adding 270 μL 37% formaldehyde) at room temperature for 10 minutes (in a total of 10mL). Formaldehyde was quenched by glycine (adding 440ul 2.5M glycine to achieve final concentration of 0.125μM), iced-cold PBS was used to wash twice. Crosslinked cells were subsequently lysed with ice cold Hi-C lysis buffer for 30 minutes at 4°C with rotation. Pelleted nuclei were digested by 375U MboI (NEB, R0147) for 2 hours at 37°C with rotation. DNA ends were filled in with biotin-dATP (Thermo, 19524016). Next, T4 DNA ligase (NEB, M0202) was added for proximity ligation for 4 hours at room temperature with rotation. Pelleted nuclei lysed with nuclear lysis buffer were sonicated with Covaris M220 (cycles/burst 200) for 240 seconds. Protein A magnetic beads were used for preclearing for 1 hour at 4°C, followed by overnight incubation with 7.5ug anti-H3K27ac antibody (Abcam, ab4729). Protein A magnetic beads were added for 2 hours to capture DNA-protein complexes. Complexes bound magnetic beads were washed followed by ChIP DNA elution. DNA was then reverse crosslinked by Proteinase K and purified with PCR purification kit. Biotin-dATP labeled DNA was used for library preparation for high-throughput sequencing as described previously by Illumina NextSeq platform (PE 75bp).

### CRISPR genome editing

CRISPR-Cas9 knockout or EBV genome editing were performed as previously described^106^. Briefly, Cas9+ Akata or Mutu I were transduced with sgRNA-expressing lentivirus and selected by puromycin (3μg/mL) or hygromycin (100μg/mL) for 7 or 14 days, respectively. EBV genomic editing efficiency was measured by indel-seq^107^. All sgRNAs were listed in the **Supplemental Table 2**.

### CRISPR interference (CRISPRi) and CRISPR activation (CRISPRa)

CRISPRi and CRISPRa were performed as previously described^108^. Briefly, EBV+ Akata cells or Mutu I cells were transduced with lentivirus expressing dCas9-KRAB or dCas9-VP64. 48 hours post-transduction, transduced cells were selected by blasticidin (5μg/mL) for 14 days to generate stable cell pools. dCas9-KRAB or dCas9-VP64 stably expressing cells were transduced with sgRNA-expressing lentivirus followed by puromycin selection for 7 days.

### Indel seq

Akata genomic DNA was harvested by QuickExtract™ DNA Extraction Solution (Biosearch Technologies), following the manufacturer’s manual. Amplicons were PCR amplified using PrimeSTAR HS DNA Polymerase (Takara) with the primers listed in **Supplemental Table 1** for Indel-seq. PCR products were purified via PCR purification kit (Qiagen) according to the manufacturer’s instructions and subject to Sanger sequencing. %indel was scored as the ratio of mutated read counts to all read counts.

### Immunoblot

Immunoblot analysis was conducted as described previously^40^. Briefly, WCL were prepared by lysing samples with 1x Lammeli buffer supplemented with β-mercaptoethanol (5%) followed by incubation for 10 minutes at 95°C. WCL were then separated by SDS-PAGE gel, transferred onto nitrocellulose membranes and blocked with 5% skim milk in TBST buffer. Primary antibodies were added for overnight incubation at 4°C. Membranes were washed with TBST buffer and incubated with secondary antibodies (Cell Signaling) for 1 hour at room temperature. SuperSignal West Chemiluminescent Substrate (Thermo Fisher Scientific) was applied for 2 minutes followed by imaging on a Li-Cor Odyssey Fc. Densitometry analysis was conducted with ImageStudioLite Odyssey software (v 5.2.5).

### RT-qPCR

RT-qPCR analysis was performed as described previously^18^. In brief, RNA was extracted with RNeasy Mini Plus Kit (Qiagen) with genomic DNA removal. M-MLV Reverse Transcriptase (Thermo Fisher Scientific) was used to synthesize cDNA. Primers for RT-qPCR are listed in **Supplemental Table 6**. Quantitative real-time PCR assays were performed with SYBR Green (Fisher Scientific). Target gene expression level was normalized to β-actin.

### EBV genome copy number quantification

EBV genome copy number quantification was performed as described previously^18^. For intracellular EBV genome copy number quantification, cells were harvested for DNA extraction using the Blood and Cell Culture DNA Minikit (Qiagen). To quantify extracellular EBV genome copy number, supernatants were collected. 20μL DNase I (10mg/mL, Promega) was used to digest unencapsidated DNA for 1 hour at 37°C. 100μL 10% SDS and 30μL proteinase K (20μg/mL, Invitrogen) were added and samples were incubated for 1 hour at 65°C. DNA was extracted by phenol-chloroform-IAA and precipitated by ethanol with sodium acetate. Purified DNA was diluted to 10ng/μL for qPCR analysis. Standard curves were generated by pHAGE-BALF5 plasmid serial dilution. Viral genome copy number was calculated using a linear regression equation.

### Flow cytometry analysis

1 × 10^6^ cells were collected and washed twice with FACS buffer (PBS with 2% FBS). Cells were stained with αgp350 antibody 72A1 for 1 hour at room temperature, washed with FACS buffer and anayzed on a BD FACSCalibur instrument. Analysis was conducted with FlowJo X software.

### Chromatin Conformation Capture (3C)

3C assays were performed as described^40^. 1 × 10^7^ cells were cross-linked in 10mL of PBS containing 10% FBS with 2% formaldehyde for 10 minutes at room temperature. Formaldehyde was quenched with glycine for 5 minutes at room temperature. Cells were pelleted, washed with ice-cold PBS and lysed with 5 mL of ice-cold Hi-C lysis buffer (10 mM Tris-HCl pH 7.5, 10 mM NaCl, 0.2% NP-40). Nuclei were collected and resuspended in 500μL of 1.2x Fastdigest restriction buffer (Thermo Fisher Scientific). 15μL of 10% SDS was added, followed by incubation for 1 hour at 37°C with shaking. A final concentration of 2% Triton X-100 was added and samples were incubated for 1 hour incubation at 37°C. 40μL of Csp6I (10U/μL, Thermo Fisher Scientific) was used for overnight digestion at 37°C overnight with shaking. 80μL of 10% SDS were added and samples were incubated at 65°C for 25 minutes with shaking. Samples were diluted 10-fold with 1.15X T4 DNA ligase reaction buffer (B0202S, New England Biolabs), and Triton X-100 was added to a final concentration of 1%. The samples were then incubated for 1 hour at 37°C with shaking at 900 rpm. Next, 100 U of T4 DNA ligase (M0202S, New England Biolabs) was added, and the reaction was carried out at 16°C for 4 hours, followed by a 30-minute incubation at room temperature. Proteinase K (300 µg, P8107S, New England Biolabs) was added, and the reaction was carried out at 65°C overnight. RNA was digested by adding 300 µg of RNase A (EN0531, Thermo Fisher) and incubating the sample for 1 hour at 37°C. As a ligation control, 10 µg of EBV bacmid was digested with 20 U of Csp6I overnight and then incubated with 20 U of T4 DNA ligase at 16°C overnight. DNA was extracted using phenol-chloroform and precipitated with ethanol. Purified DNA was then analyzed by quantitative PCR. The ΔCt method was used to analyze qPCR data, normalizing Ct values for each primer pair by setting the Ct value of 25 ng of EBV bacmid control random ligation matrix DNA to 1. All primers were listed in **Supplemental Table 7**.

### Chromatin Immunoprecipitation (ChIP)

1 × 10^7^ EBV+ Burkitt cells were cross-linked in 10mL culture media with 1% formaldehyde. Cells were lysed in ChIP lysis buffer (50mM Tris-HCl pH7.5, 10mM EDTA, 1%SDS) supplemented with protease inhibitor cocktail (Roche, cOmplete Protease Inhibitor Cocktail). Chromatin was fragmented by Bioruptor Pico sonication device (Diagenode). Fragmented chromatin was diluted and precleared with Salmon sperm DNA/Protein A agarose beads for 1 hour. Antibodies were added to pre-cleared Burkitt cell lysates for overnight incubation at 4°C. Immunocomplexes were incubated with protein A or protein G beads (25 μL) for 2 hours at 4°C. Beads were washed twice with low salt wash buffer (20mM Tris-HCl pH7.5, 150mM NaCl, 2mM EDTA, 1% Triton X-100, 0.1% SDS), high salt wash buffer (20mM Tris-HCl pH7.5, 500mM NaCl, 2mM EDTA, 1% Triton X-100, 0.1% SDS) and LiCl wash buffer (20mM Tris-HCl pH7.5, 250mM LiCl, 1mM EDTA, 1% NP-40, 1% sodium deoxycholate). DNA was eluted, reverse cross-linked by proteinase K (NEB) and purified by QIAquick PCR Purification Kit (Qiagen). Purified DNA was then subjected to qPCR analysis or high-throughput sequencing. For ChIP-seq, DNA libraries were prepared with NEBNext® Ultra™ II DNA Library Prep Kit for Illumina (NEB).

### Dual luciferase assays

Luciferase assays were conducted using the Dual-Luciferase Reporter Assay System (Promega), following the manufacturer’s instructions^109^. Briefly, Cas9+ Akata cells with Tet-inducible BZLF1 were transduced with lentivirus expressing sgRNAs against control *BXLF1* or against the *BZLF1* promoter. Transduced cells were puromycin selected. Cells were then electroporated by the Neon system (Invitrogen, same program as used for electroporation above) with 0.5μg negative control empty vector or with the *BZLF1* promoter firefly luciferase construct, together with a control *Renilla* luciferase vector (0.5μg). 24 hours post-transfection, cells were treated with doxycycline (0.5μg/mL) for 24 hours, which induces BZLF1 expression and which in turn activates the BZLF1 promoter. Cells were then lysed with Passive Lysis Buffer (Promega) for 20 min at room temperature. 20µL of lysates were added to 96-well plates, and 100µL of LAR II luciferase assay reagent (Promega) was dispensed into each well and mixed. Firefly luciferase activity was measured. 100µL of Stop and Glo Reagent (Promega) was dispensed into each well and mixed, and *Renilla* luciferase activity was measured. Luminescence was measured with a Spectramax L Microplate Reader (Molecular Devices). Firefly:*Renilla* luciferase activity ratios were calculated.

### Rapid amplification of cDNA ends (RACE)

RACE assays were conducted with SMARTer RACE 5’/3’ Kit (Takara) according to the manufacturer’s protocol. EBV+ Akata cells were treated with TPA/NaB for 24 hours. Total RNA was extracted with TRIzol Reagent (Invitrogen) followed by DNase I treatment. 1µg of total RNA was used for first-strand cDNA synthesis. For 3’-first-strand cDNA synthesis, poly(A) tails were added using Poly(A) Polymerase (Takara) prior to reverse transcription. For 5’-first strand cDNA synthesis, random primers and SMARTer II A Oligonucleotide were used for reverse transcription. Gene-specific primers (GSPs) were designed according to the manufacturer’s instructions with 15bp overlaps, using the pRACE vector. Touchdown PCR was performed with GSPs in a thermal cycler (94°C, 30 sec, 72°C, 3 min for 5 cycles; 94°C, 30 sec, 70°C, 30 s, 72°C, 3 min for 5 cycles; 94°C, 30 sec, 68°C, 30 s, 72°C, 3 min for 25 cycles and a final extension at 72°C, 2 min). RACE products were processed to In-Fusion cloning and analyzed by Sanger sequencing. Primers for RACE are listed in **Supplemental Table 8**.

### Quantification of pro-lytic and pro-latency *oriLyt* lncRNA levels

Plasmids carrying the DNA sequence of pro-lytic and pro-latency *oriLyt* lncRNAs were linearized and *in vitro* transcribed using the MAXIscript T7 Transcription Kit (Thermo) at 37°C for 1h. DNA templates were digested by TURBO DNase at 37°C for 15 min. RNA was purified by ethanol precipitation. Serial dilution of *in vitro* transcribed pro-lytic and pro-latency *oriLyt* lncRNAs were subjected to strand-specific reverse transcription. Standard curves of Ct values and input RNA copy number were generated by qPCR analysis of serial dilutions of *in vitro* transcribed lncRNAs, followed by plotting with linear regression analysis. RNA extracted from 500,000 cells were quantified by RT-qPCR with pro-lytic and pro-latency *oriLyt* lncRNAs specific primers to obtain the Ct values. Then, copy numbers of RNAs per cell from 500,000 cells were calculated based on the standard curve generated above. Corin and C12 were used at 5µM for the indicated time.

### Immunoprecipitation

5×10^7^ cells were harvested and lysed in ice cold IP lysis buffer (50mM Tris, 250mM NaCl, 0.5% NP-40 in dH_2_O) supplemented with 1mM Na3VO4, 1mM NaF and cOmplete protease inhibitor cocktail (Sigma) at 4℃ for 1h with rotation. Debris was pelleting by microcentrifugation and incubated overnight with anti-HA tag magnetic beads (Thermo) at 4℃. Beads were washed with IP lysis buffer four times and proteins were eluted with 1X Lammeli buffer supplemented with β-mercaptoethanol (5%), followed by incubation at 95°C for 10 minutes. Samples were subjected to immunoblot analysis.

### Chromatin Isolation by RNA Purification

Chromatin Isolation by RNA Purification (ChIRP) was conducted as previously described^110^. Biotinylated probes were designed using probe designer (Singlemoleculefish.com) and synthesized by IDT. Probes were labeled based on their RNA positions and separated into two pools, labeled “odd” and “even”. 2×10^7^ Akata EBV+ cells were harvested and cross-linked with 1% fresh glutaraldehyde (Sigma-Aldrich) in 10mL PBS at room temperature for 10 min. A final concentration of 0.125M glycine was used to quench the cross-linking reaction. Cell pellet was washed with 20mL chilled PBS. ChIRP lysis buffer, supplemented with cOmplete protease inhibitor cocktail, PMSF and Superase-In, was used to resuspend pellets (1mL ChIRP lysis buffer for 0.1g cell pellet). Lysates were sonicated by Bioruptor (30 sec On, 45 sec Off pulse intervals, 60 cycles) until lysates were clear. Lysates distributed between two sonication tubes were pooled together and then redistributed into two sonication tubes every 30 minutes. 5µL of lysate was extracted to analyze DNA fragment sizes. Lysates were centrifuged at 15,000g at 4°C for 10 min and combined. 10% of lysate volumes were taken for RNA and DNA inputs and kept on ice. 2mL of ChIRP hybridization buffer (50 mM Tris-HCl pH 7.0, 750 mM NaCl, 1% SDS, 1 mM EDTA, 15% formamide supplemented with cOmplete protease inhibitor cocktail, PMSF and Superase-In) was added to the lysates. Pools of 100 pmol odd or even probes against pro-lytic or pro-latency *oriLyt* lncRNA were added and mixtures were incubated at 37°C for 4h with shaking. To enrich RNA-chromatin complexes, 100 µL of streptavidin magnetic beads were washed with ChIRP lysis buffer and incubated with lysates at 37°C for 30 min with shaking. 1mL pre-warmed ChIRP wash buffer (2X sodium citrate, diluted from a 20X stock (Invitrogen, 15557044), 0.5% SDS supplemented with PMSF(1mM) was used to wash beads 5 times at 37°C with shaking. 100µL beads were removed for RNA isolation and diluted with (1 mL) of TRIZOL, followed by treatment with DNAse I (10mg/ml) for 15 minutes and then strand-specific RT-qPCR was performed. DNA was eluted from the remaining pelleted beads by incubation with RNase A (10 mg/mL), RNase H (10 U/uL) and proteinase K (40U/μL) for 30 min at 37°C. DNA was purified by incubation with phenol-chloroform (pH 7.9), ethanol precipitated, and resuspended in 50 μL. DNA samples were subjected to library preparation for Illumina NextSeq platform (PE 75bp) high-throughput sequencing. All probes were listed in the **Supplemental Table 5**.

### *In vitro* transcription and RNA pulldown assays

Plasmids templates were linearized and *in vitro* transcribed by the MAXIscript T7 Transcription Kit (Thermo) with 25% biotinylated UTP (bio-16-UTP, Thermo) and 75% unbiotinylated UTP at 37°C for 1h. Turbo DNase was used to digest DNA templates at 37°C for 15 min. Purified RNA prepared in RNA structure buffer (10mM Tris pH7.0, 100mM KCl, 10mM MgCl2 supplemented with Superase-In) was heated at 95°C for 5 min and slowly cooled to 20°C. Biotinylated RNA was incubated with streptavidin magnetic beads (50μL) at room temperature for 30 min with rotation. Nuclear extracts were prepared from 5 million Akata cells, using RNA pulldown lysis buffer (25mM Tris-HCl pH7.0, 150mM KCl, 0.5% NP-40, 1mM EDTA, 5% glycerol supplemented with cOmplete protease inhibitor cocktail and PMSF) and pre-cleared with streptavidin magnetic beads and incubated with yeast tRNA (10 mg/mL, Thermo) at 4°C for 20 min. Nuclear extracts were diluted with RIP buffer (25mM Tris-HCl pH7.0, 150mM KCl, 0.5mM DTT, 0.05% NP-40, 1mM EDTA, 5% glycerol supplemented with cOmplete protease inhibitor cocktail, PMSF and Superase-In) and incubated with 25 pmoilof biotinylated RNA at 4°C for 4h. Beads were then washed four times with high salt wash buffer (25mM Tris pH7.0, 500mM NaCl, 0.05% TritonX-100). Proteins were eluted by 1X Lammeli buffer supplemented with β-mercaptoethanol, followed by incubation at 95°C for 10 minutes.

### RNA immunoprecipitation (RIP)

RIP assays were performed using Magna RIP Kit (Millipore) according to the manufacturer’s protocol. Briefly, 2×10^7^ Akata EBV+ cells were harvested and washed with iced-cold PBS. Pelleted cells were resuspended in RIP lysis buffer supplemented with cOmplete protease inhibitors and RNase inhibitor to prepare cell lysates. Protein A/G magnetic beads (30uL, Thermo Fisher Scientific) coated with 5μg of αCTCF (Millipore) or αIgG (Millipore) were incubated with cell lysates overnight at 4 ℃. Beads were washed 4 times with RIP wash buffer for 5 min. Co-precipitated RNAs were eluted with elution buffer (provided by the Magna RIP Kit) according to the manufacturer’s instructions, purified by iodoacetic acid:phenol:chloroform (25:24:1) at pH 7.9.

### DMSMap-seq

DMS modification of EBV RNA in Akata cells were performed. Total RNA was extracted with rRNA depleted followed by amplicon-based DMS-MaPseq library generation. Briefly, Akata cells were lytically induced once they have reached a density of 10^5^ cells/ml. After 24hrs post induction cells resuspended in 5ml of complete growth medium probed with 250ul of DMS (5%) and incubated for 5 mins at 37°C. DMS reaction is quenched by adding 1.5ml (30%) of β-ME and spun down at 3,000rcf for 3 mins. Cell pellet is washed with 10ml of DPBS with 30% BME and spun down at 3,000rcf for 3mins; repeated for a total of two washes. Cell pellet is washed two more times with 10ml of DPBS and spun down at 3,000rcf for 3mins. Pellet is resuspended in 1ml of TRIzol. RNA was extracted using TRIzol’s protocol with the aqueous phase transferred to a new tube and mixed with an equal volume of 100% EtOH. Total RNA was then purified using RNA Clean and Concentrator −25 kit (Zymo). 10ug of total RNA per reaction were resuspended in 6ul of RNase-free water and used as the input for rRNA subtraction. 1μl of rRNA oligo mix (10 μg/μl) and 3 μl of hybridization buffer (200 mM NaCl, 100 mM Tris-HCl, pH 7.4) were added to each sample and incubated at 95 °C for 2 min and then the temperature was reduced by 0.1 °C/s until 45 °C were reached. Afterwards, a preheated mix of 10 μl of RNase H buffer (NEB) and 2μl thermostable RNase H (NEB) was added and incubated at 45 °C for 30 min. The RNA was cleaned with an RNA Clean and Concentrator −5 (Zymo) and eluted in 42μl water. Then, 5μl Turbo DNase buffer and 3μl Turbo DNase (Invitrogen) were added to each reaction and incubated for 20 min at 37 °C. The RNA was purified with an RNA Clean and Concentrator −5 (Zymo) and eluted in 10ul. 1ug of ribodepleted RNA was reverse transcribed using reverse primers specific for either Sense or Antisense RNA (Table 1). Primers were designed to breakdown the RNA of interest into six overlapping amplicons ranging from 150-300 nucleotides in length. For each amplicon a reverse transcription (RT) reaction was prepared by mixing 1ug of RNA, 0.5ul of Single-stranded Binding (SSB) protein, 4ul of 10mM reverse primer, 2ul of 10mM dNTPs, 0.4ul of RNase Inhibitor, 8ul of Induro RT reaction buffer (NEB), 2ul of Induro Reverse transcriptase (NEB), and completed to a final volume of 40ul with RNase-free water. Reactions were incubated at 60°C for 30 mins and then 2ul of 4M NaOH were added to the reactions and incubated for 3 mins at 95°C. RT reactions were cleaned up using a DNA Clean and Concentrator kit -5 (Zymo) and eluted in 10ul. The resulting 10ul of cDNA was PCR amplified by mixing 25ul of the Platinum SuperFi II DNA Polymerase, 2.5ul of 10mMcustom primers for each amplicon, and completed to a final volume of 50ul. Reactions went through an initial denaturation at 98°C for 30 s; 25 cycles of 98°C for 10 s, 63.6°C for 10 s, and 72°C for 20 s; and final extension at 72°C for 2mins. PCR reactions were cleaned up using a DNA Clean and Concentrator kit -5 (Zymo) and eluted in 10ul. The resulting amplicons of the same samples were pooled together, and library prepped using the NEBNext UltraTM II FS DNA Library Prep Kit for Illumina. The ≤ 100 ng input protocol was followed exactly as is except for the final indexing PCR which was performed using 25ul of the Platinum SuperFi II DNA Polymerase. Reactions went through an initial denaturation at 98°C for 30 s; 4-5 cycles of 98°C for 10 s, 65°C for 10 s, and 72°C for 60 s; and final extension at 72°C for 4mins. The resulting libraries were quantified using the Qubit 1X dsDNA HS kit and expected sizes were confirmed using E-Gel EX Agarose Gels. Samples were pooled and sequenced using a the NextSeq 1000 Sequencing System (Illumina) with 2 x 150 bp paired-end reads according to the manufacturer’s protocol.

Sequencing data were processed with SEISMIC-RNA v0.20^111^ (https://github.com/rouskinlab/seismic-rna) to compute mutation rates, graphs, Gini indexes, AUC values, and secondary structure models. Primer binding sites, reads with more than 10% fraction of mutated bases, as well as G and U bases were masked before any graph and model production. Structure models were generated using VARNA^112^ and bases were colored using normalized DMS signals. AUC values were determined for individual amplicons, Values displayed in figure belong to the amplicons that encompass the structure of interest.

### CD spectroscopy

CD spectroscopy was performed at the Harvard Medical School Center for Macromolecular Interactions on a Jasco J-1500 Spectropolarimeter (JASCO), using a 1mm optical path length quartz cuvette. A final concentration of 10µM oligos were prepared in 10mM LiCac buffer (pH 7.2) containing 1mM EDTA and 150mM LiCl or KCl. Samples were heated at 95°C for 5 min and slowly cooled down to 25 °C. Scans were conducted at an 1nm interval from 200 to 320nm at 25°C. A blank control with 10mM LiCac buffer (pH 7.2) containing 1mM EDTA and 150mM KCl was subtracted from the experimental sample data. Data were normalized to molar ellipticity.

### BG4 RIP

1×10^7^ Akata cells were washed with ice-cold PBS. Pelleted cells were lysed with RIP lysis buffer supplemented with cOmplete protease inhibitors and RNase inhibitor (1:100) at 4℃ for 20 min with rotation. Cell lysates were pre-cleared with 50μL Protein G Dynabeads (Thermo) at 4 ℃ for 1h. Protein G beads were coated with 5μg BG4 antibody (Absolute) or αIgG (Millipore) were incubated with pre-cleared cell lysates at 4℃ overnight. Beads were washed 4 times with RIP wash buffer for 5 min. Co-precipitated RNAs were isolated by elution buffer, purified by acid phenol as described above, and used for quantitative RT-PCR analysis.

### Psoralen cross-linking, nuclease digestion and RNA proximal ligation

Psoralen cross-linking followed by nuclease digestion and RNA proximal ligation assays were performed as described^77^. Briefly, Akata cells (5 million) were resuspended in pre-chilled aminomethyltrioxsalen solution (0.5 mg/ml in PBS). After incubation on ice for 15 min, cells were transferred to tissue culture plates and cross-linked with 400mJ/cm2 254nm UV light (Spectrolinker™ XL-1000 UV Crosslinker). Crosslinked cells were pelleted by centrifuging at 1,200 rpm for 5 min. RNA were extracted by TRIzol followed by isopropanol precipitation. DNase I was used to remove genomic DNA. 20µg RNA was fragmented with 1X fragmentation buffer (Thermo, AM8740) and heated at 70°C for 15 min. Fragmented RNA was purified with acid phenol:chloroform (1 mL) as above, followed by dephosphorylation by recombinant shrimp alkaline phosphatase (New England Biolabs) at 37°C for 30 min. RNA was 5’-phosphorylated by T4 PNK at 37°C for 30 min and extracted using a RNeasy Mini Kit (Qiagen) with a final elution in 25µl RNase-free H_2_O. 12µl RNA was mixed with 5 µl 10× T4 RNA ligase buffer (New England Biolabs), 3 µl T4 RNA ligase I (New England Biolabs), 2.5 µl of 100 mM ATP and 25 µl H_2_O supplemented with 2.5 µl Superase-In for overnight incubation at 16°C. M-MLV Reverse Transcriptase (Thermo Fisher Scientific) was used for cDNA synthesis. Samples without proximal ligation were used as controls for qPCR.

### Sequencing quality evaluation

All sequencing reads, including ChIP-seq, ChIRP-seq and HiChIP, were quality controlled using FastQC v0.11.9 to ensure no significant GC bias, adapters and PCR artifacts (https://www.bioinformatics.babraham.ac.uk/projects/fastqc).

### Akata EBV genome mapping

Paired-end RNA-seq reads were aligned to the Akata EBV genome using *STAR* v2.7.3a^113^ with the aligning parameters “*--outSAMprimaryFlag AllBestScore*”. ChIP-seq and ChIRP-seq reads were aligned to the Akata EBV genome using *Bowtie2* v2.5.1^114^ with the parameters *-k 1*”. PCR duplicated reads were marked and removed using *Picard* (http://broadinstitute.github.io/picard/) and *Samtools^115^*. HiChIP paired-end reads were mapped to the Akata EBV genome using HiC-Pro v2.11.4 (Servant et al., 2015) (default settings with LIGATION_SITE set as GATCGATC for MboI).

### Akata EBV visualization

For Akata EBV tracks, total reads numbers in each group were used to normalize RNA-seq, ChIP-seq and ChIRP-seq data. Normalized tracks were generated using *bedtools* (https://github.com/arq5x/bedtools2) v2.30.0 with the parameters “genomecov -bga -split -scale” together with *bedGraghToBigWig* (http://hgdownload.cse.ucsc.edu/admin/exe/linux.x86_64/bedGraphToBigWig). Tracks were visualized using circos *v0.69.8^116^* and *IGV^117^* (https://igv.org/). Significant chromatin interactions were calculated, normalized and visualized based on the analysis pipeline described^108^.

### Statistical analyses

Statistical analyses were carried out using GraphPad Prism (version 8.0; GraphPad Inc., La Jolla, CA, USA), IBM SPSS (version 23.0; IBM Corp., Armonk, NY, USA), and R (version 3.6) software for Windows. Data were obtained from at least three independent experiments and statistical significance was analyzed using a two-tailed unpaired Student’s *t*-test between two groups and by one-way analysis of variance (ANOVA) followed by a Bonferroni test for multiple comparisons. A *P*-value of <0.05 was considered statistically significant.

### Graphics and data visualization

Graphs were made with GraphPad Prism. Schematic models were made with Biorender. ChIP-seq and ChIRP data were visualized using the IGV browser.

### Resource availability

Lead contact: Ben Gewurz.

### Material availability

Materials and protocols will be shared upon request.

### Data and code availability

No new code was written for these studies. Bioinformatic programs used were commercially available and are described above.

## Supplemental figure legends

**Figure S1. Lytic induction promotes *oriLyt* and *BZLF1* promoter long-range interaction.**

A. EBV genomic H3K27ac HiChIP circos plots, as in Fig. 1B. Plots demonstrate long-range DNA interactions between H3K27Ac-marked EBV genomic regions (red arcs) in Akata Burkitt cells, mock induced (left) or induced for lytic reactivation (right) by αIgG for 24 hours (10μg/ml). Gray and orange arcs indicate short range interactions within 20 kilobases (<20kb) vs long range interactions >20 kilobases, respectively. Data are from 2 independent replicates. Genomic position of selected EBV genes, the rightward and leftward origins of lytic replication (*oriLyt^R^* and *oriLyt^L^*), the origin of plasmid replication (*oriP*) and the terminal repeats (TR) are indicated. Also shown are Akata RAD21 (orange) and CTCF (blue) ChIP-seq tracks from two replicates.
B. *OriLyt* CRISPR interference impairs EBV lytic reactivation. Immunoblots of WCL from dCas9-KRAB+ Mutu I Burkitt cells expressing the indicated sgRNAs and mock induced or induced for EBV lytic reactivation by αIgM (5μg/mL) for 4 hours.
C. Mean ± S.D. of DNase-treated extracellular EBV genome copy number from 3 independent replicates of dCas9-KRAB+ Akata cells expressing the indicated sgRNAs and mock induced or induced for EBV reactivation by αIgG crosslinking for 24 hours. *P < 0.05
D. Immunoblots of WCL from dCas9-KRAB+ Akata cells expressing control or sgRNAs targeting BILF2 and induced or induced for EBV lytic reactivation by αIgM (5μg/mL) crosslinking for 24 hours.
E. Immunoblot of WCL from dCas9-KRAB+ Akata cells expressing control or sgRNAs targeting *oriLyt^R^ or oriLyt^L^* and induced for EBV reactivation by TPA (20 ng/ml) and sodium butyrate (NaB, 2mM) treatment for 24 hours.
F. 3C assay analysis of *oriLyt* CRISPRi effects on TPA/NaB driven EBV genomic long-range interactions. Shown are the mean ± S.D. interaction frequencies, normalized by EBV bacmid levels, from 3 replicates of 3C analyses using the indicated BZLF1 anchor region (shown in blue) primer and the indicated test (T) primer regions in dCas9-KRAB+ Akata cells expressing the indicated sgRNAs and treated with TPA/NaB as in (D). P-values compare the indicated to αIgG BXLF1 control values. Blots are representative of n=3 replicates. *P < 0.05

**Figure S2. CRISPR editing of *oriLyt* or *BZLF1p* CBS compromises RAD21 cohesin loading and EBV reactivation.**

A. ChIP-qPCR analysis of Akata B cell CTCF (left) and RAD21 (right) occupancy at *oriLyt* and *BZLF1* promoter sites following mock or TPA (20 ng/ml) + NaB (2mM) treatment for 8 hours. Mean ± S.D. ChIP-qPCR values from 3 replicates.
B. Validation of the RAD21 degron system. αRAD21 immunoblot analysis of WCL from Cas9+ Akata cells expressing RAD21-FKBP12^F36V^ and control versus RAD21 sgRNA expression, as indicated.
C. Immunoblot analysis of WCL from Akata cells with stable RAD21-FKBP12^F36V^ expression, treated with DMSO vehicle versus dTAG-13 (100nM) for the indicated hours. Shown to the right are mean ± S.D. MYC/GAPDH immunoblot densitometry values from n=3 replicates.
D. Immunoblot of WCL from Akata cells with doxycycline-induced BZLF1 transgene expression, treated with dTAG-13 for 2 hours followed by doxycycline (0.5) for 24 hours.
E. Indel-seq analysis of editing activity at BZLF1 promoter (left) and *oriLyt* in Cas9+ Akata cells. Shown are the insertion/deletion (indel) percentages at *BZLF1p* vs *oriLyt* in cells expressing sgRNA targeting control *BXLF1*, a non-CBS containing (nCBS) *BZLF1p* control site, the *BZLF1p* CBS or the *oriLyt* CBS.
F. ChIP-qPCR analysis of IgG negative control versus CTCF occupancy at *BZLF1p*, *oriLyt* and *LMP2A*p in Cas9+ Akata cells expressing the sgRNA targeting control *BXLF1*, the BZLF1p nCBS control site, the *BZLF1p* CBS or the *oriLyt* CBS. Shown are mean ± S.D. % of input values from 3 replicates.
G. Analysis of CBS CRISPR editing effects on inherent *BZLF1* promoter activity. To test whether CRISPR editing of the *BZLF1p* CBS altered inherent BZLF1 promoter activity, we performed dual luciferase assays. The indicated sgRNAs were expressed in Cas9+ Akata cells and then electroporated with vectors encoding a promoterless firefly luciferase control or with a *BZLF1* promoter sequence upstream of firefly luciferase, together with control renilla luciferase. To test *BZLF1* promoter activity, we induced expression of a BZLF1 transgene by addition of doxycycline (0.5μg/ml) for 24hr (BZLF1 binds to the BZLF1 promoter and upregulates its activity^118^). Shown are mean renilla-normalized firefly luciferase values ± S.D. from 3 independent replicates.
H. Analysis of control *BXLF1* or *BILF2* CBS CRISPR editing effects on EBV maintenance of latency+. Immunoblot analysis of WCL from Cas9+ Akata cells expressing *BXLF1* or *BILF2*-CBS targeting sgRNAs.
I. RT-qPCR analysis of BZLF1 mRNA expression from Cas9+ Akata cells with sgRNAs targeting control *BXLF1* or *BILF2* CBS. Shown are mean ± S.D. BZLF1 mRNA levels from n=3 replicates, with the mean BXLF1 control cell level set to 1. P-values compare the indicated with BXLF1 control levels. Immunoblots are representative of n=3 replicates. *P<0.05, **P<0.01

**Figure S3. Key pro-lytic lncRNA roles in oriLyt:BZLF1 promoter interaction and lytic reactivation.**

A. Analysis of αIgG crosslinking effects on *oriLyt* H3K27ac levels. Mean ± S.D. values from 3 replicates of ChIP-qPCR analysis of H3K27ac *oriLyt* levels in Akata cells, mock stimulated or stimulated by αIgG for 2 hours.
B. ChIP-qPCR analysis of RNA polymerase II (Pol II), Pol II serine 2 phosphorylation (S2P) and Pol II serine 5 phosphorylation (S5P) levels at *oriLyt* in Akata cells mock stimulated or stimulated by αIgG for 2 hours. Mean ± S.D. values from 3 replicates.
C. Strand-specific PROSEQ analysis ^57^ of nascent *oriLyt^R^* region transcripts in Mutu I cells treated with TPA/NaB for the indicated hours. The black bar depicts the *oriLyt^R^*region, the grey bar indicates highly shared oriLyt^R^ and oriLyt^L^ sequence.
D. Agarose gel analysis of 5’- and 3’-RACE PCR products, using three sets of gene specific primers (#1-3). At right are representative Sanger DNA sequencing traces of sense and antisense oriLyt lncRNA obtained by 5’-RACE.
E. Validation of lncRNA copy number assay. Shown are mean Ct values from qPCR analysis of antisense (left) or sense (right) *oriLyt* lncRNA copy number, using serial dilutions of *in vitro* transcribed lncRNAs. Standard curves were generated by linear regression analysis.
F. Mean ± S.D. sense and antisense *oriLyt* lncRNA levels in unstimulated Akata cells. Shown are average lncRNA copy number, calculated according to Ct value and normalized by molecular weight and input cell number.
G. Mean ± S.D. *oriLyt* sense and antisense lncRNA levels from 3 RT-PCR replicates performed on Akata cells that were stimulated by αIgG for the indicated times.
H. Mean ± S.D. *oriLyt* sense versus antisense lncRNA levels from 3 RT-PCR replicates performed on Akata cells stimulated by TPA/NaB for the indicated times.
I. Validation of shRNA mediated lncRNA knockdown. Mean ± S.D. *oriLyt* antisense versus sense lncRNA levels from n=3 RT-PCR replicates of Akata cells expressing control shRNA or shRNAs targeting the antisense lncRNA (left) or sense lncRNA (right) and stimulated by αIgG crosslinking for 24 hours.
J. Effects of *oriLyt* sense lncRNA knockdown on EBV lytic reactivation. Immunoblots of WCL from Akata cells that expressed control shRNA or *oriLyt* sense lncRNA targeting shRNA and following αIgG crosslinking for 24hours.
K. Effects of *oriLyt anti*sense lncRNA knockdown on EBV lytic reactivation. Immunoblots of WCL from Akata cells with control shRNA or *oriLyt* antisense lncRNA targeting shRNAs and following αIgG crosslinking for 24hours.
L. Validation of *oriLyt* lncRNA knockdown by locked nucleic acids. Mean ± S.D. *oriLyt* pro-latency (antisense, left) and pro-lytic (sense, right) lncRNA levels from 3 replicates of strand-specific RT-qPCR analysis of Akata cells expressing control LNA versus LNA targeting either the pro-latency or pro-lytic lncRNAs.
M. 3C assay analysis of long-range interactions between the BZLF1 promoter and EBV genomic regions. Mean ± S.D. values from 3 replicates of 3C assays performed on Akata cells with negative control shRNA or shRNAs targeting the *oriLyt* pro-lytic lncRNA. Cells were mock stimulated or stimulated by TPA/NaB treatment for 24 hours, as indicated. 3C assay frequencies were normalized by EBV bacmid input values. P-value cross-compares values from cells with shRNA control vs shRNA targeting the pro-lytic lncRNAs. Blots are representative of n=3 replicates. *p < 0.05

**Figure S4. Lytic-induced MYC degradation unloads LSD1 in *oriLyt*.**

A. Effects of MYC overexpression on *oriLyt* total Pol II and Pol II Ser2P occupancy. Mean ± S.D. values from 3 ChIP-qPCR replicates, using control IgG, Pol II (left) or Pol II phosphoserine 2 specific antibodies (right) and *oriLyt* primers in Akata cells expressing control GFP or *MYC* cDNAs. Cells were mock stimulated or stimulated with αIgG crosslinking for 2 hours.
B. Analysis of MIZ-1 depletion effects on EBV reactivation. Immunoblot analysis of WCL from Cas9+ Akata that expressed control or three independent MIZ-1 targeting sgRNAs. Blots are representative of n=3 replicates.
C. Relative protein levels of MYC, LSD1, ZNF217 and CoREST in P3HR-1 Burkitt cells (orange) or in P3HR-1 with conditional BZLF1 and BRLF1 immediate early proteins, mock triggered or triggered for reactivation for the indicated hours^58^. P3HR-1 with conditional BZLF1/BRFL1 were FACsorted for abortive gp350- (orange) versus fully lytic gp350+ cells (green) prior to proteomic analysis.
D. Analysis of TPA/NaB effects on EBV+ Burkitt MYC levels. Mean ± S.D. values from 3 RT-qPCR replicates of GAPDH-normalized MYC mRNA levels in Akata cells that were stimulated by TPA/NaB (TPA 20 ng/ml, NaB 2mM) for the indicated times.
E. Analysis of CRISPR *oriLyt* EBox editing on LSD1 occupancy. Mean ± S.D. values from n=3 ChIP-qPCR replicates of using control or α-LSD1 antibodies and primers specific for *oriLyt* in BZLF1-knockout Akata cells that expressed control or oriLyt Ebox targeting sgRNAs^14^.
F. Analysis of FACT antagonist CBL0137 effects on *oriLyt* LSD1 occupancy. Mean ± S.D. values from 3 ChIP-qPCR replicates, using control or α-LSD1 antibodies and primers specific for *oriLyt* in BZLF1-knockout Akata cells that were treated with vehicle control or with CBL0137 (5μM) for 24 hours. CBL0137 treatment depletes B cell MYC expression ^40^.
G. Validation of LSD1, ZNF217 or CoREST depletion effects on EBV reactivation. Immunoblot analysis of WCL from Akata cells expressing control sgRNA or *LSD1, ZNF217* or *CoREST* targeting sgRNAs. Blot is representative of n=3 replicates.
H. Analysis of LSD1, ZNF217 or CoREST depletion effects on *oriLyt* pro-lytic lncRNA expression. Mean ± S.D. values from 3 replicates of strand-specific analysis of pro-lytic *oriLyt* expression level in Cas9+ Akata cells expressing the indicated sgRNAs. For cross-comparison, qPCR analysis of GAPDH normalized BZLF1 mRNA levels are shown.

*P<0.05, **P<0.01.

**Figure S5. *oriLyt* transcription supports cohesin-mediated *oriLyt:BZLF1* looping.**

A. CRISPRa *oriLyt* pro-lytic lncRNA upregulation. Mean ± S.D. GAPDH-normalized *oriLyt* pro-lytic lncRNA levels from 3 qPCR replicates using dCas9-VP64+ Akata cells expressing control vs pro-lytic lncRNA promoter targeting sgRNAs.
B. O*riLyt* pro-lytic lncRNA expression upregulates *oriLyt* RNA Pol II occupancy. Mean ± S.D. values from 3 ChIP-qPCR analysis replicates of RNA Pol II (left) or Pol II phosphoserine 2 (S2P, a mark of elongating Pol II, right) *oriLyt* occupancy in dCas9-VP64+ Akata cells expressing control (grey boxes) or pro-lytic lncRNA promoter targeting sgRNA (red and blue boxes).
C. Analysis of CRISPRa upregulation of *oriLyt* pro-lytic lncRNA effects on EBV lytic reactivation. Immunoblot analysis of WCL from dCas9-VP64+ Mutu I cells expressing the indicated BXLF1 control versus independent oriLyt pro-lytic lncRNA targeting sgRNA for 5 days.
D. Opposing EBV reactivation effects of CRISPRa upregulated *oriLyt* pro-latency vs pro-lytic lncRNAs. Immunoblot analysis of WCL from dCas9-VP64+ Mutu I cells expressing control BXLF1 sgRNA or sgRNAs targeting the promoters of *oriLyt* pro-latency vs pro-lytic lncRNAs. Cells were stimulated by αIgG for 24 hours, as indicated.
E. CRISPRa *oriLyt* pro-lytic lncRNA upregulation induces *oriLyt*:*BZLF1* promoter interactions. Mean ± S.D. 3C assay values from 3 replicates performed on dCas9-VP64+ Akata cells that expressed control sgRNA vs sgRNA targeting the *oriLyt* pro-lytic lncRNA promoter. 3C assay frequencies were normalized by bacmid input values. *BZLF1* anchor primer (blue), *oriLyt^R^*(red) and test (T) primer regions are depicted in the schematic at top.
F. CDK9 inhibition effects on Pol II phosphorylation and on αIgG induced BZLF1 expression. Immunoblot of WCL from Akata cells treated with vehicle or with CDK9 antagonist NVP2 (0.25nM, 0.5nM and 1nM) and/or by αIgG (10μg/mL) for 24 hours.
G. NVP2 effects on *oriLyt* pro-lytic lncRNA levels. Mean ± S.D. GAPDH normalized *oriLyt* pro-lytic lncRNA qPCR levels from 3 replicates of Akata cells as in (F).
H. NVP2 effects on *BZLF1* promoter looping. Mean ± S.D. 3C assay values from 3 replicates, performed on Akata cells mock stimulated or stimulated by αIgG in the absence or presence of NVP2, as in (F).
I. NVP2 effects on *BZLF1* promoter CTCF and RAD21 loading. ChIP-qPCR analysis of CTCF and RAD21 *BZLF1* promoter occupancy in Akata cells mock stimulated or stimulated by αIgG crosslinking, in the absence or presence of NVP2, as in (F).

Immunoblots are representative of n=3 independent replicates. *p < 0.05, **p < 0.01, ***p<0.001

**Figure S6. A pro-lytic lncRNA encoded RNA G4 quadruplex recruits CTCF to oriLyt.**

A. Cross-comparison of *oriLyt* lncRNA ChIRP-seq and CTCF, LSD1 and ZNF217 ChIP-seq. EBV genomic *oriLyt* pro-lytic and pro-latency lncRNAs ChIRP-seq tracks (top) juxtaposed with CTCF and ZNF217 ChIP-seq tracks (bottom). ChIRP-seq and CTCF ChIP-seq were generated from Akata cells at 24 hours post-αIgG crosslinking. ZNF217 ChIP-seq were performed on unstimulated Akata cells^15^. *OriLyt* region zoomed in tracks are shown below.
B. Exogenous *oriLyt* pro-lytic lncRNA overexpression effects on BZLF1 induction. Mean ± S.D. values from 3 qPCR replicates of GAPDH-normalized BZLF1 levels in Akata cells at 24 hours post-electroporation of pcDNA3.1(-) empty vector control versus pcDNA3.1 (-) expressing the *oriLyt* pro-lytic lncRNA and αIgG crosslinked for 24 hours.
C. Exogenous *oriLyt* pro-lytic lncRNA overexpression effects on lytic protein expression. Immunoblots of WCL from cells as in (B).
D. Pro-lytic lncRNA knowdown effects on *oriLyt* enhancer epigenetic marks. Mean ± S.D. values from 3 replicates of H3K4me1 or H3K27ac ChIP followed by *oriLyt* qPCR using Akata cells that expressed negative control shRNA or *oriLyt* pro-lytic lncRNA targeting shRNAs and that were mock stimulated or stimulated by αIgG crosslinking for 4h.
E. TERRA RNA CD spectra. As a positive control for a lncRNA with a known rG4, 10µM TERRA was incubated in 10mM LiCac buffer (pH 7.2) containing 150mM KCl and analyzed by CD. Characteristic negative and positive peaks were observed at ∼240nm and 265nm, respectively, indicative of the presence of a G4 structure.
F. *Detection of the oriLyt* pro-lytic lncRNA rG4 *in vivo*. Mean ± S.D. values from 3 replicates of RNA immunoprecipitation (RIP) qPCR analysis of pro-lytic lncRNA or negative control EBER2 levels. The monoclonal antibody BG4 is specific for G4. RIP was performed on extracts of Akata cells that were mock simulated, stimulated by αIgG (10μg/ml) crosslinking or by TPA (20ng/mL) plus NaB (2mM) for 8h. RIP IgG control levels from mock stimulated cells were set to 1.

**Figure S7. RNA G4 structure is important for EBV reactivation.**

A. The oriLyt pro-lytic lncRNA contains an rG4 structure. CD spectra analysis of 10µM *oriLyt* pro-lytic lncRNA resuspended in 10mM LiCac buffer (pH 7.2) containing 150mM KCl with the indicated TMPyP4 concentrations. TMPyP4 binds to and disrupts rG4^76^. Reduction of the 265nm peak magnitude with increasing TMPyP4 concentrations is indicative of a G4 structure.
B. TMPyP4 inhibits EBV reactivation. Immunoblot analysis of WCL from Akata cells that were treated with vehicle or the indicated TMPyP4 concentrations and mock-stimulated or stimulated by aIgG for 24 hours. Blots are representative of n=3 replicates.
C. TMPyP4 reduces *oriLyt* CTCF occupancy. Mean ± S.D. values from 3 replicates of IgG control vs αCTCF ChIP followed by qPCR for *oriLyt*, using Akata cells that were mock stimulated or stimulated by αIgG for 4 hours ± TMPyP4 (50µM), as indicated.
D. TMPyP4 inhibits *oriLyt:BZLF1* promoter looping following αIgG crosslinking. Mean ± S.D. values from 3 replicates of 3C analysis, using a BZLF1 promoter anchor primer and test (T) primers to the indicated EBV genomic regions. Akata cells were mock stimulated or stimulated by αIgG for 24 hours ± TMPyP4 (50µM). 3C assay frequencies were normalized by bacmid input values.
E. Analysis of *oriLyt* pro-lytic and pro-latency lncRNA association following EBV reactivation. RT-PCR SYBR signals from proximity ligation assays (PLA) of *oriLyt* pro-lytic and pro-latency lncRNAs from three independent replicates. PLA was performed on lysates of Akata cells that were transfected with scrambled LNA or *oriLyt* pro-latency lncRNA targeting LNA and following αIgG crosslinking for 2 hours. Shown also is a negative control, in which no enzyme was added. Samples were psoralen crosslinked prior to PLA. RT-qPCR was conducted to detect interactions between pro-lytic and pro-latency *oriLyt* lncRNAs. *p < 0.05

## Supplemental Information.

**Supplemental Table 1. List of PCR primers used in this study.**

**Supplemental Table 2. List of sgRNAs for CRISPR editing, CRISPRi and CRISPRa in this study.**

**Supplemental Table 3. List of shRNAs used in this study.**

**Supplemental Table 4. List of locked nucleic acid (LNA) used in this study.**

**Supplemental Table 5. List of CHIRP probes used in this study.**

**Supplemental Table 6. List of qPCR primers used in this study.**

**Supplemental Table 7. List of oligonucleotides for 3C assays in this study.**

**Supplemental Table 8. List of oligonucleotides for RACE assays in this study.**

## Notes

### Competing Interest Statement

The authors have declared no competing interest.

## References

1. Munz, C. (2025). Epstein-Barr virus pathogenesis and emerging control strategies. Nat Rev Microbiol 23, 667–679. 10.1038/s41579-025-01181-y.

2. Chiu, Y.F., Ponlachantra, K., and Sugden, B. (2024). How Epstein Barr Virus Causes Lymphomas. Viruses 16. 10.3390/v16111744.

3. Farrell, P.J. (2019). Epstein-Barr Virus and Cancer. Annu Rev Pathol 14, 29–53. 10.1146/annurev-pathmechdis-012418-013023.

4. Damania, B., Kenney, S.C., and Raab-Traub, N. (2022). Epstein-Barr virus: Biology and clinical disease. Cell 185, 3652–3670. 10.1016/j.cell.2022.08.026.

5. Murata, T., Okuno, Y., Sato, Y., Watanabe, T., and Kimura, H. (2020). Oncogenesis of CAEBV revealed: Intragenic deletions in the viral genome and leaky expression of lytic genes. Rev Med Virol 30, e2095. 10.1002/rmv.2095.

6. Gru, A.A., Haverkos, B.H., Freud, A.G., Hastings, J., Nowacki, N.B., Barrionuevo, C., Vigil, C.E., Rochford, R., Natkunam, Y., Baiocchi, R.A., and Porcu, P. (2015). The Epstein-Barr Virus (EBV) in T Cell and NK Cell Lymphomas: Time for a Reassessment. Curr Hematol Malig Rep 10, 456–467. 10.1007/s11899-015-0292-z.

7. Robinson, W.H., Younis, S., Love, Z.Z., Steinman, L., and Lanz, T.V. (2024). Epstein-Barr virus as a potentiator of autoimmune diseases. Nat Rev Rheumatol 20, 729–740. 10.1038/s41584-024-01167-9.

8. SoRelle, E.D., and Luftig, M.A. (2025). Multiple sclerosis and infection: history, EBV, and the search for mechanism. Microbiol Mol Biol Rev 89, e0011923. 10.1128/mmbr.00119-23.

9. Guo, R., and Gewurz, B.E. (2022). Epigenetic control of the Epstein-Barr lifecycle. Curr Opin Virol 52, 78–88. 10.1016/j.coviro.2021.11.013.

10. Chakravorty, A., Sugden, B., and Johannsen, E.C. (2019). An Epigenetic Journey: Epstein-Barr Virus Transcribes Chromatinized and Subsequently Unchromatinized Templates during Its Lytic Cycle. J Virol 93. 10.1128/JVI.02247-18.

11. Buschle, A., and Hammerschmidt, W. (2020). Epigenetic lifestyle of Epstein-Barr virus. Semin Immunopathol 42, 131–142. 10.1007/s00281-020-00792-2.

12. Kenney, S.C., and Mertz, J.E. (2014). Regulation of the latent-lytic switch in Epstein-Barr virus. Semin Cancer Biol 26, 60–68. 10.1016/j.semcancer.2014.01.002.

13. Thorley-Lawson, D.A. (2015). EBV Persistence--Introducing the Virus. Curr Top Microbiol Immunol 390, 151–209. 10.1007/978-3-319-22822-8_8.

14. Guo, R., Jiang, C., Zhang, Y., Govande, A., Trudeau, S.J., Chen, F., Fry, C.J., Puri, R., Wolinsky, E., Schineller, M., et al. (2020). MYC Controls the Epstein-Barr Virus Lytic Switch. Mol Cell 78, 653–669 e658. 10.1016/j.molcel.2020.03.025.

15. Liao, Y., Yan, J., Kong, I., Li, Z., Ding, W., Clark, S., Giulino-Roth, L., and Gewurz, B.E. (2025). The Histone Demethylase LSD1/ZNF217/CoREST Complex is a Major Restriction Factor of Epstein-Barr Virus Lytic Reactivation. Res Sq. 10.21203/rs.3.rs-5649616/v1.

16. Guo, R., Zhang, Y., Teng, M., Jiang, C., Schineller, M., Zhao, B., Doench, J.G., O’Reilly, R.J., Cesarman, E., Giulino-Roth, L., and Gewurz, B.E. (2020). DNA methylation enzymes and PRC1 restrict B-cell Epstein-Barr virus oncoprotein expression. Nat Microbiol 5, 1051–1063. 10.1038/s41564-020-0724-y.

17. Zhang, Y., Jiang, C., Trudeau, S.J., Narita, Y., Zhao, B., Teng, M., Guo, R., and Gewurz, B.E. (2020). Histone Loaders CAF1 and HIRA Restrict Epstein-Barr Virus B-Cell Lytic Reactivation. mBio 11. 10.1128/mBio.01063-20.

18. Murray-Nerger, L.A., Lozano, C., Burton, E.M., Liao, Y., Ungerleider, N.A., Guo, R., and Gewurz, B.E. (2024). The nucleic acid binding protein SFPQ represses EBV lytic reactivation by promoting histone H1 expression. Nat Commun 15, 4156. 10.1038/s41467-024-48333-x.

19. Countryman, J., and Miller, G. (1985). Activation of expression of latent Epstein-Barr herpesvirus after gene transfer with a small cloned subfragment of heterogeneous viral DNA. Proc Natl Acad Sci U S A 82, 4085–4089. 10.1073/pnas.82.12.4085.

20. Feederle, R., Kost, M., Baumann, M., Janz, A., Drouet, E., Hammerschmidt, W., and Delecluse, H.J. (2000). The Epstein-Barr virus lytic program is controlled by the co-operative functions of two transactivators. EMBO J 19, 3080–3089. 10.1093/emboj/19.12.3080.

21. Ali, A., Ohashi, M., Casco, A., Djavadian, R., Eichelberg, M., Kenney, S.C., and Johannsen, E. (2022). Rta is the principal activator of Epstein-Barr virus epithelial lytic transcription. PLoS Pathog 18, e1010886. 10.1371/journal.ppat.1010886.

22. Wille, C.K., Nawandar, D.M., Panfil, A.R., Ko, M.M., Hagemeier, S.R., and Kenney, S.C. (2013). Viral genome methylation differentially affects the ability of BZLF1 versus BRLF1 to activate Epstein-Barr virus lytic gene expression and viral replication. J Virol 87, 935–950. 10.1128/JVI.01790-12.

23. Ragoczy, T., and Miller, G. (1999). Role of the epstein-barr virus RTA protein in activation of distinct classes of viral lytic cycle genes. J Virol 73, 9858–9866. 10.1128/JVI.73.12.9858-9866.1999.

24. Zalani, S., Holley-Guthrie, E., and Kenney, S. (1996). Epstein-Barr viral latency is disrupted by the immediate-early BRLF1 protein through a cell-specific mechanism. Proc Natl Acad Sci U S A 93, 9194–9199. 10.1073/pnas.93.17.9194.

25. Meng, Q., Hagemeier, S.R., Fingeroth, J.D., Gershburg, E., Pagano, J.S., and Kenney, S.C. (2010). The Epstein-Barr virus (EBV)-encoded protein kinase, EBV-PK, but not the thymidine kinase (EBV-TK), is required for ganciclovir and acyclovir inhibition of lytic viral production. J Virol 84, 4534–4542. 10.1128/JVI.02487-09.

26. Murata, T. (2018). Encyclopedia of EBV-Encoded Lytic Genes: An Update. Adv Exp Med Biol 1045, 395–412. 10.1007/978-981-10-7230-7_18.

27. Hammerschmidt, W. (2015). The Epigenetic Life Cycle of Epstein-Barr Virus. Curr Top Microbiol Immunol 390, 103–117. 10.1007/978-3-319-22822-8_6.

28. Djavadian, R., Chiu, Y.F., and Johannsen, E. (2016). An Epstein-Barr Virus-Encoded Protein Complex Requires an Origin of Lytic Replication In Cis to Mediate Late Gene Transcription. PLoS Pathog 12, e1005718. 10.1371/journal.ppat.1005718.

29. Hammerschmidt, W., and Sugden, B. (1988). Identification and characterization of oriLyt, a lytic origin of DNA replication of Epstein-Barr virus. Cell 55, 427–433. 10.1016/0092-8674(88)90028-1.

30. Xue, S.A., and Griffin, B.E. (2007). Complexities associated with expression of Epstein-Barr virus (EBV) lytic origins of DNA replication. Nucleic Acids Res 35, 3391–3406. 10.1093/nar/gkm170.

31. Westphal, E.M., Blackstock, W., Feng, W., Israel, B., and Kenney, S.C. (2000). Activation of lytic Epstein-Barr virus (EBV) infection by radiation and sodium butyrate in vitro and in vivo: a potential method for treating EBV-positive malignancies. Cancer Res 60, 5781–5788.

32. Li, H., Lee, C.Y., and Delecluse, H.J. (2025). Epstein-Barr virus lytic replication and cancer. Curr Opin Virol 70, 101438. 10.1016/j.coviro.2024.101438.

33. Ma, S.D., Yu, X., Mertz, J.E., Gumperz, J.E., Reinheim, E., Zhou, Y., Tang, W., Burlingham, W.J., Gulley, M.L., and Kenney, S.C. (2012). An Epstein-Barr Virus (EBV) mutant with enhanced BZLF1 expression causes lymphomas with abortive lytic EBV infection in a humanized mouse model. J Virol 86, 7976–7987. 10.1128/JVI.00770-12.

34. Ma, S.D., Hegde, S., Young, K.H., Sullivan, R., Rajesh, D., Zhou, Y., Jankowska-Gan, E., Burlingham, W.J., Sun, X., Gulley, M.L., et al. (2011). A new model of Epstein-Barr virus infection reveals an important role for early lytic viral protein expression in the development of lymphomas. J Virol 85, 165–177. 10.1128/JVI.01512-10.

35. Okuno, Y., Murata, T., Sato, Y., Muramatsu, H., Ito, Y., Watanabe, T., Okuno, T., Murakami, N., Yoshida, K., Sawada, A., et al. (2019). Defective Epstein-Barr virus in chronic active infection and haematological malignancy. Nat Microbiol 4, 404–413. 10.1038/s41564-018-0334-0.

36. SoRelle, E.D., Haynes, L.E., Willard, K.A., Chang, B., Ch’ng, J., Christofk, H., and Luftig, M.A. (2024). Epstein-Barr virus reactivation induces divergent abortive, reprogrammed, and host shutoff states by lytic progression. PLoS Pathog 20, e1012341. 10.1371/journal.ppat.1012341.

37. Mumbach, M.R., Rubin, A.J., Flynn, R.A., Dai, C., Khavari, P.A., Greenleaf, W.J., and Chang, H.Y. (2016). HiChIP: efficient and sensitive analysis of protein-directed genome architecture. Nat Methods 13, 919–922. 10.1038/nmeth.3999.

38. Nabet, B., Roberts, J.M., Buckley, D.L., Paulk, J., Dastjerdi, S., Yang, A., Leggett, A.L., Erb, M.A., Lawlor, M.A., Souza, A., et al. (2018). The dTAG system for immediate and target-specific protein degradation. Nat Chem Biol 14, 431–441. 10.1038/s41589-018-0021-8.

39. Qi, L.S., Larson, M.H., Gilbert, L.A., Doudna, J.A., Weissman, J.S., Arkin, A.P., and Lim, W.A. (2013). Repurposing CRISPR as an RNA-guided platform for sequence-specific control of gene expression. Cell 152, 1173–1183. 10.1016/j.cell.2013.02.022.

40. Guo, R., Jiang, C., Zhang, Y., Govande, A., Trudeau, S.J., Chen, F., Fry, C.J., Puri, R., Wolinsky, E., Schineller, M., et al. (2020). MYC Controls the Epstein-Barr Virus Lytic Switch. Mol Cell 78, 653–669.e658. 10.1016/j.molcel.2020.03.025.

41. Peters, J.M., Tedeschi, A., and Schmitz, J. (2008). The cohesin complex and its roles in chromosome biology. Genes Dev 22, 3089–3114. 10.1101/gad.1724308.

42. Yatskevich, S., Rhodes, J., and Nasmyth, K. (2019). Organization of Chromosomal DNA by SMC Complexes. Annu Rev Genet 53, 445–482. 10.1146/annurev-genet-112618-043633.

43. Kim, Y., Shi, Z., Zhang, H., Finkelstein, I.J., and Yu, H. (2019). Human cohesin compacts DNA by loop extrusion. Science 366, 1345–1349. 10.1126/science.aaz4475.

44. Davidson, I.F., Bauer, B., Goetz, D., Tang, W., Wutz, G., and Peters, J.M. (2019). DNA loop extrusion by human cohesin. Science 366, 1338–1345. 10.1126/science.aaz3418.

45. Hashimoto, H., Wang, D., Horton, J.R., Zhang, X., Corces, V.G., and Cheng, X. (2017). Structural Basis for the Versatile and Methylation-Dependent Binding of CTCF to DNA. Mol Cell 66, 711–720 e713. 10.1016/j.molcel.2017.05.004.

46. Yin, M., Wang, J., Wang, M., Li, X., Zhang, M., Wu, Q., and Wang, Y. (2017). Molecular mechanism of directional CTCF recognition of a diverse range of genomic sites. Cell Res 27, 1365–1377. 10.1038/cr.2017.131.

47. Nichols, M.H., and Corces, V.G. (2015). A CTCF Code for 3D Genome Architecture. Cell 162, 703–705. 10.1016/j.cell.2015.07.053.

48. Croce, C.M., Erikson, J., ar-Rushdi, A., Aden, D., and Nishikura, K. (1984). Translocated c-myc oncogene of Burkitt lymphoma is transcribed in plasma cells and repressed in lymphoblastoid cells. Proc Natl Acad Sci U S A 81, 3170–3174. 10.1073/pnas.81.10.3170.

49. Andersson, R., and Sandelin, A. (2020). Determinants of enhancer and promoter activities of regulatory elements. Nat Rev Genet 21, 71–87. 10.1038/s41576-019-0173-8.

50. Kaikkonen, M.U., Spann, N.J., Heinz, S., Romanoski, C.E., Allison, K.A., Stender, J.D., Chun, H.B., Tough, D.F., Prinjha, R.K., Benner, C., and Glass, C.K. (2013). Remodeling of the enhancer landscape during macrophage activation is coupled to enhancer transcription. Mol Cell 51, 310–325. 10.1016/j.molcel.2013.07.010.

51. Lam, M.T., Cho, H., Lesch, H.P., Gosselin, D., Heinz, S., Tanaka-Oishi, Y., Benner, C., Kaikkonen, M.U., Kim, A.S., Kosaka, M., et al. (2013). Rev-Erbs repress macrophage gene expression by inhibiting enhancer-directed transcription. Nature 498, 511–515. 10.1038/nature12209.

52. Li, W., Notani, D., Ma, Q., Tanasa, B., Nunez, E., Chen, A.Y., Merkurjev, D., Zhang, J., Ohgi, K., Song, X., et al. (2013). Functional roles of enhancer RNAs for oestrogen-dependent transcriptional activation. Nature 498, 516–520. 10.1038/nature12210.

53. Melo, C.A., Drost, J., Wijchers, P.J., van de Werken, H., de Wit, E., Oude Vrielink, J.A., Elkon, R., Melo, S.A., Léveillé, N., Kalluri, R., et al. (2013). eRNAs are required for p53-dependent enhancer activity and gene transcription. Mol Cell 49, 524–535. 10.1016/j.molcel.2012.11.021.

54. Xiang, J.F., Yin, Q.F., Chen, T., Zhang, Y., Zhang, X.O., Wu, Z., Zhang, S., Wang, H.B., Ge, J., Lu, X., et al. (2014). Human colorectal cancer-specific CCAT1-L lncRNA regulates long-range chromatin interactions at the MYC locus. Cell Res 24, 513–531. 10.1038/cr.2014.35.

55. Schaukowitch, K., Joo, J.Y., Liu, X., Watts, J.K., Martinez, C., and Kim, T.K. (2014). Enhancer RNA facilitates NELF release from immediate early genes. Mol Cell 56, 29–42. 10.1016/j.molcel.2014.08.023.

56. O’Grady, T., Cao, S., Strong, M.J., Concha, M., Wang, X., Splinter Bondurant, S., Adams, M., Baddoo, M., Srivastav, S.K., Lin, Z., et al. (2014). Global bidirectional transcription of the Epstein-Barr virus genome during reactivation. J Virol 88, 1604–1616. 10.1128/JVI.02989-13.

57. Dunn, L.E.M., Lu, F., Su, C., Lieberman, P.M., and Baines, J.D. (2023). Reactivation of Epstein-Barr Virus from Latency Involves Increased RNA Polymerase Activity at CTCF Binding Sites on the Viral Genome. J Virol 97, e0189422. 10.1128/jvi.01894-22.

58. Ersing, I., Nobre, L., Wang, L.W., Soday, L., Ma, Y., Paulo, J.A., Narita, Y., Ashbaugh, C.W., Jiang, C., Grayson, N.E., et al. (2017). A Temporal Proteomic Map of Epstein-Barr Virus Lytic Replication in B Cells. Cell Rep 19, 1479–1493. 10.1016/j.celrep.2017.04.062.

59. Wiese, K.E., Walz, S., von Eyss, B., Wolf, E., Athineos, D., Sansom, O., and Eilers, M. (2013). The role of MIZ-1 in MYC-dependent tumorigenesis. Cold Spring Harb Perspect Med 3, a014290. 10.1101/cshperspect.a014290.

60. Zhang, L., Li, J., Xu, H., Shao, X., Fu, L., Hou, Y., Hao, C., Li, W., Joshi, K., Wei, W., et al. (2020). Myc-Miz1 signaling promotes self-renewal of leukemia stem cells by repressing Cebpα and Cebpδ. Blood 135, 1133–1145. 10.1182/blood.2019001863.

61. Kalkat, M., Resetca, D., Lourenco, C., Chan, P.K., Wei, Y., Shiah, Y.J., Vitkin, N., Tong, Y., Sunnerhagen, M., Done, S.J., et al. (2018). MYC Protein Interactome Profiling Reveals Functionally Distinct Regions that Cooperate to Drive Tumorigenesis. Mol Cell 72, 836–848.e837. 10.1016/j.molcel.2018.09.031.

62. Kalin, J.H., Wu, M., Gomez, A.V., Song, Y., Das, J., Hayward, D., Adejola, N., Wu, M., Panova, I., Chung, H.J., et al. (2018). Targeting the CoREST complex with dual histone deacetylase and demethylase inhibitors. Nat Commun 9, 53. 10.1038/s41467-017-02242-4.

63. Sorna, V., Theisen, E.R., Stephens, B., Warner, S.L., Bearss, D.J., Vankayalapati, H., and Sharma, S. (2013). High-throughput virtual screening identifies novel N’-(1-phenylethylidene)-benzohydrazides as potent, specific, and reversible LSD1 inhibitors. J Med Chem 56, 9496–9508. 10.1021/jm400870h.

64. Olson, C.M., Jiang, B., Erb, M.A., Liang, Y., Doctor, Z.M., Zhang, Z., Zhang, T., Kwiatkowski, N., Boukhali, M., Green, J.L., et al. (2018). Pharmacological perturbation of CDK9 using selective CDK9 inhibition or degradation. Nat Chem Biol 14, 163–170. 10.1038/nchembio.2538.

65. Chu, C., Zhang, Q.C., da Rocha, S.T., Flynn, R.A., Bharadwaj, M., Calabrese, J.M., Magnuson, T., Heard, E., and Chang, H.Y. (2015). Systematic discovery of Xist RNA binding proteins. Cell 161, 404–416. 10.1016/j.cell.2015.03.025.

66. Bose, D.A., Donahue, G., Reinberg, D., Shiekhattar, R., Bonasio, R., and Berger, S.L. (2017). RNA Binding to CBP Stimulates Histone Acetylation and Transcription. Cell 168, 135–149.e122. 10.1016/j.cell.2016.12.020.

67. Huang, Z., Liang, N., Goñi, S., Damdimopoulos, A., Wang, C., Ballaire, R., Jager, J., Niskanen, H., Han, H., Jakobsson, T., et al. (2021). The corepressors GPS2 and SMRT control enhancer and silencer remodeling via eRNA transcription during inflammatory activation of macrophages. Mol Cell 81, 953–968.e959. 10.1016/j.molcel.2020.12.040.

68. Saldaña-Meyer, R., Rodriguez-Hernaez, J., Escobar, T., Nishana, M., Jácome-López, K., Nora, E.P., Bruneau, B.G., Tsirigos, A., Furlan-Magaril, M., Skok, J., and Reinberg, D. (2019). RNA Interactions Are Essential for CTCF-Mediated Genome Organization. Mol Cell 76, 412–422.e415. 10.1016/j.molcel.2019.08.015.

69. Zubradt, M., Gupta, P., Persad, S., Lambowitz, A.M., Weissman, J.S., and Rouskin, S. (2017). DMS-MaPseq for genome-wide or targeted RNA structure probing in vivo. Nat Methods 14, 75–82. 10.1038/nmeth.4057.

70. Luige, J., Armaos, A., Tartaglia, G.G., and Ørom, U.A.V. (2024). Predicting nuclear G-quadruplex RNA-binding proteins with roles in transcription and phase separation. Nat Commun 15, 2585. 10.1038/s41467-024-46731-9.

71. Lee, Y.W., Weissbein, U., Blum, R., and Lee, J.T. (2024). G-quadruplex folding in Xist RNA antagonizes PRC2 activity for stepwise regulation of X chromosome inactivation. Mol Cell 84, 1870–1885 e1879. 10.1016/j.molcel.2024.04.015.

72. Takahama, K., and Oyoshi, T. (2013). Specific binding of modified RGG domain in TLS/FUS to G-quadruplex RNA: tyrosines in RGG domain recognize 2’-OH of the riboses of loops in G-quadruplex. J Am Chem Soc 135, 18016–18019. 10.1021/ja4086929.

73. Martadinata, H., and Phan, A.T. (2013). Structure of human telomeric RNA (TERRA): stacking of two G-quadruplex blocks in K(+) solution. Biochemistry 52, 2176–2183. 10.1021/bi301606u.

74. Biffi, G., Tannahill, D., McCafferty, J., and Balasubramanian, S. (2013). Quantitative visualization of DNA G-quadruplex structures in human cells. Nat Chem 5, 182–186. 10.1038/nchem.1548.

75. Morris, M.J., Wingate, K.L., Silwal, J., Leeper, T.C., and Basu, S. (2012). The porphyrin TmPyP4 unfolds the extremely stable G-quadruplex in MT3-MMP mRNA and alleviates its repressive effect to enhance translation in eukaryotic cells. Nucleic Acids Res 40, 4137–4145. 10.1093/nar/gkr1308.

76. Zamiri, B., Reddy, K., Macgregor, R.B., Jr., and Pearson, C.E. (2014). TMPyP4 porphyrin distorts RNA G-quadruplex structures of the disease-associated r(GGGGCC)n repeat of the C9orf72 gene and blocks interaction of RNA-binding proteins. J Biol Chem 289, 4653–4659. 10.1074/jbc.C113.502336.

77. Culbertson, B., Garcia, K., Markett, D., Asgharian, H., Chen, L., Fish, L., Navickas, A., Yu, J., Woo, B., Nanda, A.S., et al. (2023). A sense-antisense RNA interaction promotes breast cancer metastasis via regulation of NQO1 expression. Nat Cancer 4, 682–698. 10.1038/s43018-023-00554-7.

78. Lu, Z., Gong, J., and Zhang, Q.C. (2018). PARIS: Psoralen Analysis of RNA Interactions and Structures with High Throughput and Resolution. Methods Mol Biol 1649, 59–84. 10.1007/978-1-4939-7213-5_4.

79. Babcock, G.J., Decker, L.L., Freeman, R.B., and Thorley-Lawson, D.A. (1999). Epstein-barr virus-infected resting memory B cells, not proliferating lymphoblasts, accumulate in the peripheral blood of immunosuppressed patients. J Exp Med 190, 567–576. 10.1084/jem.190.4.567.

80. Schepers, A., Pich, D., and Hammerschmidt, W. (1996). Activation of oriLyt, the lytic origin of DNA replication of Epstein-Barr virus, by BZLF1. Virology 220, 367–376. 10.1006/viro.1996.0325.

81. Pombo, A., and Dillon, N. (2015). Three-dimensional genome architecture: players and mechanisms. Nat Rev Mol Cell Biol 16, 245–257. 10.1038/nrm3965.

82. Lieberman, P.M. (2025). Transcriptional regulation of gene modules in Epstein-Barr virus. Transcription, 1–21. 10.1080/21541264.2025.2562704.

83. Caruso, L.B., Maestri, D., and Tempera, I. (2023). Three-Dimensional Chromatin Structure of the EBV Genome: A Crucial Factor in Viral Infection. Viruses 15. 10.3390/v15051088.

84. Lee, S.H., Kim, K.D., Cho, M., Huh, S., An, S.H., Seo, D., Kang, K., Lee, M., Tanizawa, H., Jung, I., et al. (2023). Characterization of a new CCCTC-binding factor binding site as a dual regulator of Epstein-Barr virus latent infection. PLoS Pathog 19, e1011078. 10.1371/journal.ppat.1011078.

85. Hughes, D.J., Marendy, E.M., Dickerson, C.A., Yetming, K.D., Sample, C.E., and Sample, J.T. (2012). Contributions of CTCF and DNA methyltransferases DNMT1 and DNMT3B to Epstein-Barr virus restricted latency. J Virol 86, 1034– 1045. 10.1128/JVI.05923-11.

86. Takacs, M., Banati, F., Koroknai, A., Segesdi, J., Salamon, D., Wolf, H., Niller, H.H., and Minarovits, J. (2010). Epigenetic regulation of latent Epstein-Barr virus promoters. Biochim Biophys Acta 1799, 228–235. 10.1016/j.bbagrm.2009.10.005.

87. Liu, C., Kudo, T., Ye, X., and Gascoigne, K. (2023). Cell-to-cell variability in Myc dynamics drives transcriptional heterogeneity in cancer cells. Cell Rep 42, 112401. 10.1016/j.celrep.2023.112401.

88. Padovan-Merhar, O., Nair, G.P., Biaesch, A.G., Mayer, A., Scarfone, S., Foley, S.W., Wu, A.R., Churchman, L.S., Singh, A., and Raj, A. (2015). Single mammalian cells compensate for differences in cellular volume and DNA copy number through independent global transcriptional mechanisms. Mol Cell 58, 339–352. 10.1016/j.molcel.2015.03.005.

89. Dang, C.V. (2012). MYC on the path to cancer. Cell 149, 22–35. 10.1016/j.cell.2012.03.003.

90. Koganti, S., Clark, C., Zhi, J., Li, X., Chen, E.I., Chakrabortty, S., Hill, E.R., and Bhaduri-McIntosh, S. (2015). Cellular STAT3 functions via PCBP2 to restrain Epstein-Barr Virus lytic activation in B lymphocytes. J Virol 89, 5002–5011. 10.1128/JVI.00121-15.

91. Hill, E.R., Koganti, S., Zhi, J., Megyola, C., Freeman, A.F., Palendira, U., Tangye, S.G., Farrell, P.J., and Bhaduri-McIntosh, S. (2013). Signal transducer and activator of transcription 3 limits Epstein-Barr virus lytic activation in B lymphocytes. J Virol 87, 11438–11446. 10.1128/JVI.01762-13.

92. Daigle, D., Megyola, C., El-Guindy, A., Gradoville, L., Tuck, D., Miller, G., and Bhaduri-McIntosh, S. (2010). Upregulation of STAT3 marks Burkitt lymphoma cells refractory to Epstein-Barr virus lytic cycle induction by HDAC inhibitors. J Virol 84, 993–1004. 10.1128/JVI.01745-09.

93. Nakahashi, H., Kieffer Kwon, K.R., Resch, W., Vian, L., Dose, M., Stavreva, D., Hakim, O., Pruett, N., Nelson, S., Yamane, A., et al. (2013). A genome-wide map of CTCF multivalency redefines the CTCF code. Cell Rep 3, 1678–1689. 10.1016/j.celrep.2013.04.024.

94. Ohlsson, R., Lobanenkov, V., and Klenova, E. (2010). Does CTCF mediate between nuclear organization and gene expression? Bioessays 32, 37–50. 10.1002/bies.200900118.

95. Wulfridge, P., Yan, Q., Rell, N., Doherty, J., Jacobson, S., Offley, S., Deliard, S., Feng, K., Phillips-Cremins, J.E., Gardini, A., and Sarma, K. (2023). G-quadruplexes associated with R-loops promote CTCF binding. Mol Cell 83, 3064–3079.e3065. 10.1016/j.molcel.2023.07.009.

96. Lyu, K., Chow, E.Y., Mou, X., Chan, T.F., and Kwok, C.K. (2021). RNA G-quadruplexes (rG4s): genomics and biological functions. Nucleic Acids Res 49, 5426–5450. 10.1093/nar/gkab187.

97. Bjornevik, K., Munz, C., Cohen, J.I., and Ascherio, A. (2023). Epstein-Barr virus as a leading cause of multiple sclerosis: mechanisms and implications. Nat Rev Neurol 19, 160–171. 10.1038/s41582-023-00775-5.

98. Soldan, S., Su, C., Monaco, M.C., Brown, N., Clauze, A., Andrada, F., Feder, A., Planet, P., Kossenkov, A., Schaffer, D., et al. (2023). Unstable EBV latency drives inflammation in multiple sclerosis patient derived spontaneous B cells. Res Sq. 10.21203/rs.3.rs-2398872/v1.

99. ElAbd, H., Pesesky, M., Innocenti, G., Chung, B.K., Mahdy, A.K.H., Kriukova, V., Kulsvehagen, L., Strobbe, D., Stuhler, C., Mayr, G., et al. (2025). T and B cell responses against Epstein-Barr virus in primary sclerosing cholangitis. Nat Med 31, 2306–2316. 10.1038/s41591-025-03692-w.

100. Xu, X., Zhu, N., Zheng, J., Peng, Y., Zeng, M.S., Deng, K., Duan, C., and Yuan, Y. (2024). EBV abortive lytic cycle promotes nasopharyngeal carcinoma progression through recruiting monocytes and regulating their directed differentiation. PLoS Pathog 20, e1011934. 10.1371/journal.ppat.1011934.

101. Munz, C. (2020). Tumor Microenvironment Conditioning by Abortive Lytic Replication of Oncogenic gamma-Herpesviruses. Adv Exp Med Biol 1225, 127–135. 10.1007/978-3-030-35727-6_9.

102. Tsai, M.H., Raykova, A., Klinke, O., Bernhardt, K., Gartner, K., Leung, C.S., Geletneky, K., Sertel, S., Munz, C., Feederle, R., and Delecluse, H.J. (2013). Spontaneous lytic replication and epitheliotropism define an Epstein-Barr virus strain found in carcinomas. Cell Rep 5, 458–470. 10.1016/j.celrep.2013.09.012.

103. Perrine, S.P., Hermine, O., Small, T., Suarez, F., O’Reilly, R., Boulad, F., Fingeroth, J., Askin, M., Levy, A., Mentzer, S.J., et al. (2007). A phase 1/2 trial of arginine butyrate and ganciclovir in patients with Epstein-Barr virus-associated lymphoid malignancies. Blood 109, 2571–2578. 10.1182/blood-2006-01-024703.

104. Haverkos, B., Alpdogan, O., Baiocchi, R., Brammer, J.E., Feldman, T.A., Capra, M., Brem, E.A., Nair, S., Scheinberg, P., Pereira, J., et al. (2023). Targeted therapy with nanatinostat and valganciclovir in recurrent EBV-positive lymphoid malignancies: a phase 1b/2 study. Blood Adv 7, 6339–6350. 10.1182/bloodadvances.2023010330.

105. Wang, C., Liu, X., Liang, J., Narita, Y., Ding, W., Li, D., Zhang, L., Wang, H., Leong, M.M.L., Hou, I., et al. (2023). A DNA tumor virus globally reprograms host 3D genome architecture to achieve immortal growth. Nat Commun 14, 1598. 10.1038/s41467-023-37347-6.

106. Greenfeld, H., Takasaki, K., Walsh, M.J., Ersing, I., Bernhardt, K., Ma, Y., Fu, B., Ashbaugh, C.W., Cabo, J., Mollo, S.B., et al. (2015). TRAF1 Coordinates Polyubiquitin Signaling to Enhance Epstein-Barr Virus LMP1-Mediated Growth and Survival Pathway Activation. PLoS Pathog 11, e1004890. 10.1371/journal.ppat.1004890.

107. Wang, Z., Guo, R., Trudeau, S.J., Wolinsky, E., Ast, T., Liang, J.H., Jiang, C., Ma, Y., Teng, M., Mootha, V.K., and Gewurz, B.E. (2021). CYB561A3 is the key lysosomal iron reductase required for Burkitt B-cell growth and survival. Blood 138, 2216–2230. 10.1182/blood.2021011079.

108. Ding, W., Wang, C., Narita, Y., Wang, H., Leong, M.M.L., Huang, A., Liao, Y., Liu, X., Okuno, Y., Kimura, H., et al. (2022). The Epstein-Barr Virus Enhancer Interaction Landscapes in Virus-Associated Cancer Cell Lines. J Virol 96, e0073922. 10.1128/jvi.00739-22.

109. Lupey-Green, L.N., Moquin, S.A., Martin, K.A., McDevitt, S.M., Hulse, M., Caruso, L.B., Pomerantz, R.T., Miranda, J.L., and Tempera, I. (2017). PARP1 restricts Epstein Barr Virus lytic reactivation by binding the BZLF1 promoter. Virology 507, 220–230. 10.1016/j.virol.2017.04.006.

110. Chu, C., Qu, K., Zhong, F.L., Artandi, S.E., and Chang, H.Y. (2011). Genomic maps of long noncoding RNA occupancy reveal principles of RNA-chromatin interactions. Mol Cell 44, 667–678. 10.1016/j.molcel.2011.08.027.

111. Allan, M.F., Aruda, J., Plung, J.S., Grote, S.L., Martin des Taillades, Y.J., de Lajarte, A.A., Bathe, M., and Rouskin, S. (2024). Discovery and Quantification of Long-Range RNA Base Pairs in Coronavirus Genomes with SEARCH-MaP and SEISMIC-RNA. bioRxiv. 10.1101/2024.04.29.591762.

112. Darty, K., Denise, A., and Ponty, Y. (2009). VARNA: Interactive drawing and editing of the RNA secondary structure. Bioinformatics 25, 1974–1975. 10.1093/bioinformatics/btp250.

113. Dobin, A., Davis, C.A., Schlesinger, F., Drenkow, J., Zaleski, C., Jha, S., Batut, P., Chaisson, M., and Gingeras, T.R. (2013). STAR: ultrafast universal RNA-seq aligner. Bioinformatics 29, 15–21. 10.1093/bioinformatics/bts635.

114. Langmead, B., and Salzberg, S.L. (2012). Fast gapped-read alignment with Bowtie 2. Nat Methods 9, 357–359. 10.1038/nmeth.1923.

115. Li, H., Handsaker, B., Wysoker, A., Fennell, T., Ruan, J., Homer, N., Marth, G., Abecasis, G., Durbin, R., and Genome Project Data Processing, S. (2009). The Sequence Alignment/Map format and SAMtools. Bioinformatics 25, 2078–2079. 10.1093/bioinformatics/btp352.

116. Krzywinski, M., Schein, J., Birol, I., Connors, J., Gascoyne, R., Horsman, D., Jones, S.J., and Marra, M.A. (2009). Circos: an information aesthetic for comparative genomics. Genome Res 19, 1639–1645. 10.1101/gr.092759.109.

117. Robinson, J.T., Thorvaldsdottir, H., Winckler, W., Guttman, M., Lander, E.S., Getz, G., and Mesirov, J.P. (2011). Integrative genomics viewer. Nat Biotechnol 29, 24–26. 10.1038/nbt.1754.

118. Yin, Q., Jupiter, K., and Flemington, E.K. (2004). The Epstein-Barr virus transactivator Zta binds to its own promoter and is required for full promoter activity during anti-Ig and TGF-beta1 mediated reactivation. Virology 327, 134–143. 10.1016/j.virol.2004.06.026.

119. Ma, Y., Walsh, M.J., Bernhardt, K., Ashbaugh, C.W., Trudeau, S.J., Ashbaugh, I.Y., Jiang, S., Jiang, C., Zhao, B., Root, D.E., et al. (2017). CRISPR/Cas9 Screens Reveal Epstein-Barr Virus-Transformed B Cell Host Dependency Factors. Cell Host Microbe 21, 580–591 e587. 10.1016/j.chom.2017.04.005.

120. Kalkat, M., Resetca, D., Lourenco, C., Chan, P.K., Wei, Y., Shiah, Y.J., Vitkin, N., Tong, Y., Sunnerhagen, M., Done, S.J., et al. (2018). MYC Protein Interactome Profiling Reveals Functionally Distinct Regions that Cooperate to Drive Tumorigenesis. Mol Cell 72, 836–848 e837. 10.1016/j.molcel.2018.09.031.

